# Structure of TFIIH/Rad4-Rad23-Rad33 in damaged DNA opening in Nucleotide Excision Repair

**DOI:** 10.1101/2021.01.14.426755

**Authors:** Trevor van Eeuwen, Yoonjung Shim, Hee Jong Kim, Tingting Zhao, Shrabani Basu, Benjamin A. Garcia, Craig Kaplan, Jung-Hyun Min, Kenji Murakami

**Affiliations:** Department of Biochemistry and Biophysics, Perelman School of Medicine, University of Pennsylvania, Philadelphia, PA 19104, U.S.A.; Biochemistry and Molecular Biophysics Graduate Group, Perelman School of Medicine, University of Pennsylvania, Philadelphia, Pennsylvania 19104, USA; Department of Biological Sciences, University of Pittsburgh, Pittsburgh, Pennsylvania 15260, US; Epigenetics Institute, Department of Biochemistry and Biophysics, Perelman School of Medicine, University of Pennsylvania, Philadelphia, Pennsylvania 19104, USA; Department of Chemistry and Biochemistry, Baylor University, Waco, Texas 76798, USA; Penn Center for Genome Integrity, Perelman School of Medicine, University of Pennsylvania, Philadelphia, PA 19104, U.S.A.

## Abstract

The versatile nucleotide excision repair (NER) pathway initiates as the XPC-RAD23B-CETN2 complex first recognizes DNA lesions from the genomic DNA and recruits the general transcription factor complex, TFIIH, for subsequent lesion verification. Here, we present a cryo-EM structure of an NER initiation complex containing Rad4-Rad23-Rad33 (yeast homologue of XPC-RAD23B-CETN2) and 7-subunit core TFIIH assembled on a carcinogen-DNA adduct lesion at 3.9–9.2 Å resolution. A ~30-bp DNA duplex could be mapped as it straddles between Rad4 and the Ssl2 (XPB) subunit of TFIIH on the 3’ and 5’ side of the lesion, respectively. The simultaneous binding with Rad4 and TFIIH was permitted by an unwinding of DNA at the lesion. Translocation coupled with torque generation by Ssl2 would extend the DNA opening at the lesion and deliver the damaged strand to Rad3 (XPD) in an unwound form suitable for subsequent lesion scanning and verification.

## Introduction

Genomic DNA is continuously being damaged by various endogenous and exogenous genotoxic agents. If left unrepaired, DNA lesions can impair cellular functions and lead to mutations that cause various disorders such as cancers (reviewed in ^1–3^). Among various DNA damage repair pathways, nucleotide excision repair (NER) is highly conserved among eukaryotes and serves to repair a wide variety of environmentally induced lesions, including intra-strand crosslinks induced by the sunlight (ultraviolet irradiation) or chemical crosslinkers (e.g., cisplatin), and various bulky adducts induced by environmental carcinogens present in fossil fuel combustion, cigarette smoke, cooked meat, *etc.* ^4^.

Impaired NER can decrease cell survival after UV exposure in yeast and other cell lines ^5^. In humans, mutations in key NER proteins underlie genetic disorders such as xeroderma pigmentosum (XP), Cockayne syndrome and trichothiodystrophy ^6^. Patients with XP are hypersensitive to UV light and exhibit >1000-fold increase in the risk of skin cancers including melanomas and squamous cell carcinomas. Cockayne syndromes results in premature aging with neurological abnormalities.

NER lesions can be recognized via two distinct sub-pathways, either in global genome NER (GG-NER) or in transcription-coupled NER (TC-NER) (reviewed in ^7,8^). In TC-NER, lesions are recognized when RNA polymerase II (Pol II) stalls at a lesion during transcription elongation, while in GG-NER, the XPC protein in complex with RAD23B and centrin 2 (CETN2) functions as the main sensor that recognize diverse NER lesions. Lesion-bound XPC recruits the general transcription factor IIH (TFIIH), a conserved general transcription factor required for both transcription initiation and nucleotide excision repair (NER) ^9–13^, which then scans the damaged strand and verifies the presence of a bulky lesion, aided by XPA and RPA ^14,15^. Subsequently, the lesion-containing single-stranded DNA (24-32 nucleotides) is excised by XPF-ERCC1 and XPG endonucleases on the 5’ and 3’-sides of the lesion; the resulting gap is restored by the repair synthesis and nick sealing by DNA polymerases and ligases, respectively.

The key to the remarkable versatility of GG-NER lies in its unique 2-step initiation process involving XPC and TFIIH, that bypasses the need to rely on any specific structure of a lesion ^16,17^. As a first step, XPC locates a damaged site by sensing local DNA destabilization/distortion induced by a lesion, rather than the lesion structure itself ^18–21^. Crystallographic studies of Rad4-Rad23, yeast homologues of XPC-RAD23B provides structural insights into such an indirect recognition mechanism ^22–24^. Rad4 involves four consecutive domains of the protein: the transglutaminase domain (TGD) and the β-hairpin domain 1 (BHD1) bind to a ~11-bp undamaged segment of DNA, whereas BHD2 and BHD3 together bind to a 4-bp segment including the damage. This binding results in ‘open’ DNA conformation where the damaged site is locally unwound and two nucleotide pairs harboring the lesion are flipped out of the DNA duplex, stabilized by a β-hairpin from BHD3 filling the gap in the DNA. Notably, BHD2/BHD3 selectively bind to the flipped-out nucleotides from the undamaged strand and does not make direct contacts with the damaged nucleotides, thereby enabling indirect readout of various lesions regardless of the specific structures. Further kinetic and single-molecule studies have revealed that Rad4 performs fast diffusional search while probing the DNA structural integrity that involves untwisting ^24–26^.

As a second step of the NER initiation process, the lesion-bound XPC recruits TFIIH and TFIIH verifies the presence of a genuine NER lesion (a bulky damage) through the function of the XPD(Rad3) helicase ^27,28^. For instance, a 2- to 3-bp mismatched site can be specifically bound to XPC(Rad4), but is not repaired by NER, due to failure in this verification step ^16,17^. The 10-subunit holo-TFIIH consists of a 7-subunit core complex (hereafter coreTFIIH) and a 3-subunit Cdk-activating kinase module (CAK; TFIIK in yeast) ^29,30^. The core TFIIH contains two helicase-family ATPase subunits (XPB(Ssl2)and XPD(Rad3) in human(yeast)) and 5 nonenzymatic subunits (p62(Tfb1), p52(Tfb2), p44(Ssl1), p34(Tfb4) and p8(Tfb5)). One ATPase XPB(Ssl2) shows preference for double-stranded DNA (dsDNA) over single-stranded DNA (ssDNA) and is active as an ATP-dependent DNA translocase in both transcription and NER; the other ATPase subunit, XPD(Rad3) prefers ssDNA over dsDNA and is enzymatically active as a 5’-3’ helicase only in NER ^31–35^. The CAK module functions to phosphorylate RNA polymerase II (Pol II) and is thus critical for mRNA transcription and processing ^36,37^. However, it is not required for NER reaction *in vitro* ^38^ and is shown to dissociate from TFIIH upon UV irradiation in human cells ^39^. Structural studies of TFIIH have been a challenge due to its size and its compositional and conformational heterogeneities; available TFIIH structures have so far been limited to a form engaged in transcription initiation together with Pol II in human and in yeast ^40–42^ and a more recent structure of human TFIIH in complex with DNA and XPA for NER ^43^. Higher resolution structures of TFIIH including CAK have been determined only for human TFIIH ^44,45^. Thus, the structural mechanism of how TFIIH is recruited to lesions by XPC and further propels NER initiation has been a mystery.

Here we determined the structure of yeast TFIIH/Rad4-Rad23-Rad33 bound to DNA containing a single carcinogen-DNA adduct (2-acetylaminofluorene) using cryo-electron microscopy, chemical cross-linking and mass spectrometry. A ~30 bp stretch of DNA could be mapped along with the protein complex in two distinct segments with respect to the lesion: the dsDNA at the 5’ side of the DNA lesion was bound to Ssl2, similarly as observed in the transcription initiation complex, while the duplex at the 3’ side of the lesion was bound to the TGD domain of Rad4 near Tfb1 of TFIIH. In between, the DNA was locally (~3 bp) unwound at the lesion. This unwinding enables not only the simultaneous binding with Ssl2 and Rad4 but also subsequent NER bubble formation by the translocase activity of Ssl2. The DNA was, however, not yet engaged with Rad3, indicating that the ensuing lesion verification would involve additional steps of conformational changes. Altogether, our study illustrates how Rad4(XPC) recruits TFIIH to damaged DNA in 3D structural models for the first time and provides the groundwork to visualize how the NER initiation is carried out in multiple, coordinated yet distinct steps.

## Results

### Assembly of TFIIH/Rad4-Rad23-Rad33/DNA

To assemble a TFIIH/Rad4-Rad23-Rad33/DNA complex (hereafter TFIIH/Rad4-23-33/DNA) suitable for structural study, the 7-subunit core TFIIH was isolated from yeast as previously published ^46^ with slight modifications. Full-length Rad4-Rad23 complex and Rad33 were obtained by overexpression in insect cells and bacteria, respectively. To identify a suitable DNA substrate for structural studies, we employed gel electrophoresis mobility shift assays (EMSA) with a series of dsDNA containing a 2-acetylaminofluorene adduct (AAF-dG) as an NER lesion (Figures 1A-B and S1A-C). In addition to a 29-bp DNA fragment (−14 to +14, nucleotide numbering follows the 5’-3’-direction on the damaged strand with respect to the AAF-dG at position +0), which is sufficient for assembly of the Rad4-Rad23-DNA ^22,23^, we prepared DNA fragments extending ~10 bp on either end (−24/+15, –15/+24), or on both ends (−24/+24). Two DNA constructs including the region from –24 to +15 (−24/+15, –24/+24) supported a stable co-complex formation including both TFIIH and Rad4-Rad23, whereas others (−14/+14 and –15/+24) bound to either TFIIH or Rad4-Rad23 but not to both. The 10-bp dsDNA extension could also be replaced with 10-nt ssDNA without affecting the binding (see – 24/+24 vs. –24*/+24, Figure 1B). The complexes formed in the presence of ATP (4 mM) as well as its non-hydrolysable analog, AMP-PNP (Figure 1B). Additionally, the presence of Rad33, the yeast homologue of centrin 2 (CETN2) did not impair the complex assembly (Figure S1C).

**Figure 1.**
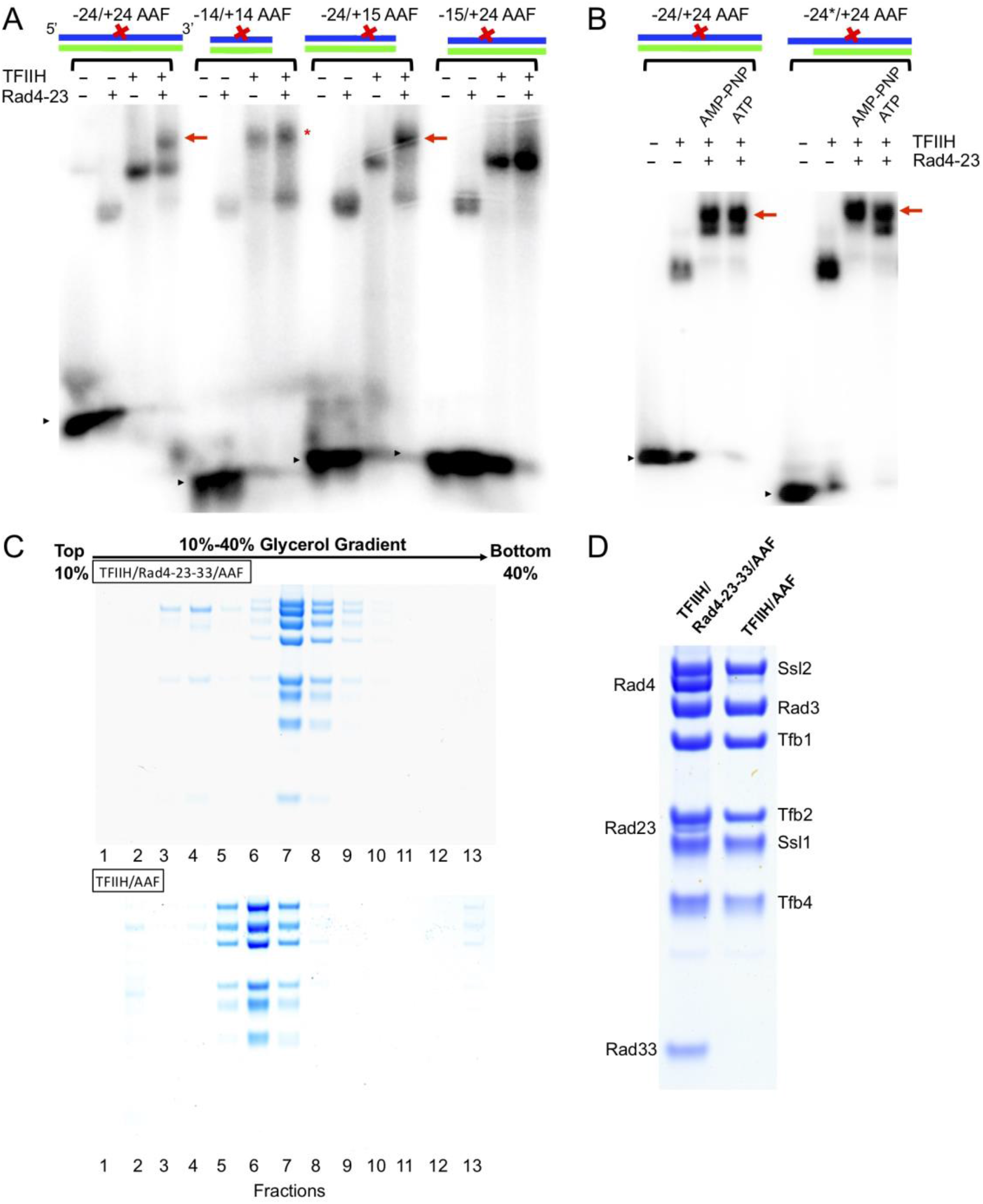
Assembly of core TFIIH and Rad4-Rad23-Rad33 on AAF-DNA. **(A)** EMSA of TFIIH and/or Rad4-Rad23 binding to various AAF-DNA constructs (schematic above) in the presence of AMP-PNP. Rad4-Rad23 and TFIIH formed complexes with −24/+24 or −24/+15 AAF-DNA (red arrow) but not with −14/+14 or −15/+24 AAF-DNA. Black arrow heads indicate unbound, free AAF-DNA. Asterisk (*) indicates non-stoichimetric TFIIH-DNA complexes lacking Rad4-Rad23. **(B)** EMSA of TFIIH with Rad4-Rad23 on −24/+24 or −24*/+24 AAF showing the complex formation (red arrow) in the presence of AMP-PNP or ATP. −24*/+24 AAF contains 10-nucleotide (nt) single-stranded overhang on the 5’ side of the DNA whereas −24/+24 AAF is fully duplexed. **(C)** −24*/+24 AAF was combined with TFIIH, Rad4-Rad23, and Rad33 (upper panel) or with TFIIH alone (lower panel) then subjected to glycerol gradient centrifugation in the presence of AMP-PNP. SDS-PAGE analysis of the fractions showed that stoichiometric amounts of TFIIH, Rad4-Rad23, and Rad33 were present in a single peak in slightly higher glycerol density (upper panel) than that of the complex with only TFIIH and DNA (lower panel), indicating the formation of a TFIIH/Rad4-Rad23-Rad33/DNA complex. Note that Rad23 is poorly stained by Coomassie. **(D)** SDS-PAGE of TFIIH/Rad4-23-33/DNA (left) and TFIIH-DNA (right) isolated by gradient sedimentation.

Following these results, we next reconstituted the TFIIH/Rad4-23-33/DNA complex on a preparative scale and sedimented on a 10–40% glycerol gradient in the presence of AMP-PNP (Figures 1C-D). Slightly excess Rad4-Rad23-Rad33 was combined with TFIIH and –24*/+24 AAF-DNA, such that free Rad4-Rad23-Rad33 appeared in lower density fractions (fractions 3-4 in upper panel of Figure 1C), while the complex containing equimolar amounts of TFIIH and Rad4-Rad23-Rad33 appeared in a single peak of higher density (fractions 7-8 in upper panel of Figure 1C). The same homogeneous complex was obtained in the presence of ATP (Figure S1E), consistent with the EMSA results (Figure 1B).

A supporting evidence that the resulting complex represents a stable entity specifically assembled on the DNA, rather than a mixture of non-specifically bound factors, came from factor challenge experiments (Figures S1D-H). When purified Rad2 (yeast homologues of XPG) was added in ~6-fold molar excess over the TFIIH/Rad4-23-33/DNA complex followed by the glycerol gradient sedimentation, ~50% of TFIIH bound Rad2, and dissociated Rad4-Rad23-Rad33 (fractions 8-11 in middle panel of Figure S1G) in the presence of ATP but not AMP-PNP (upper panel of Figure S1G). These results corroborate previous biochemical studies with HeLa nuclear extract suggesting that XPG fails to stably associate with TFIIH-XPC without ATP, and that XPG displaces XPC from TFIIH/DNA when added to TFIIH-XPC in the presence of ATP ^47,48^.

### TFIIH/Rad4-Rad23-Rad33 Cross-linking Mass Spectrometry

Further support for the specific assembly of the TFIIH/Rad4-23-33/DNA complex was obtained by cross-linking mass spectrometry (XL-MS) (Figure 2, Table S2). TFIIH/Rad4-Rad23-Rad33 assembled with the –24/+24 AAF-DNA was reacted with MS-cleavable cross-linker disuccinimidyl dibutyric urea (DSBU), and the cross-links were identified by mass spectrometry followed by computational search. A total of 333 cross-links, comprising 257 within TFIIH, 56 within Rad4-Rad23-Rad33, and 20 between TFIIH and Rad4-Rad23-Rad33, were identified. Observed cross-links between TFIIH subunits were in good agreement with those previously obtained from highly purified TFIIH ^49,50^. False positive rate of our XL-MS method was first assessed by comparison to the model of TFIIH obtained at high resolution from Map 1 (see below). Of the 257 cross-links identified here, 182 cross-links could be mapped onto corresponding residues in the model while the other 75 were on unstructured flexible loops and thus were not mapped; 18 crosslinks were between residues more than 40 Å apart (the maximum distance of cross-linking) in the structure of TFIIH, corresponding to a violation rate of 7% (Figure 2B). Of the 20 cross-links between Rad4-23-33 and TFIIH, 18 cross-links were mapped on the PH domain and BSD2 domain of Tfb1, and the C-terminal region of Ssl2, suggesting that Tfb1 and Ssl2 of TFIIH make the primary contacts with Rad4-Rad23-Rad33. Our results are completely consistent with a prevailing human NER model that the XPC protein interacts with TFIIH through the p62 and XPB subunits, human homologues of Tfb1 and Ssl2, respectively ^51,52^, and further extended the model when combined with cryo-EM data (see below). Moreover, intra-subunit cross-links within TFIIH in the TFIIH/Rad4-23-33/DNA complex significantly differed from those in holo-TFIIH containing TFIIK (Kin28-Ccl1-Tfb3) (Figures S2A-B) (e.g., there were more intra-subunit cross-links within Tfb2 (K171 – K495, S80 – K506, indicated by red dashed lines in Figure 3C), and less cross-links within Ssl2). These changes in XL-MS are in good agreement with a structural arrangement of TFIIH itself upon binding Rad4-Rad23-Rad33-DNA and/or dissociation of TFIIK by cryo-EM analysis (see below).

**Figure 2.**
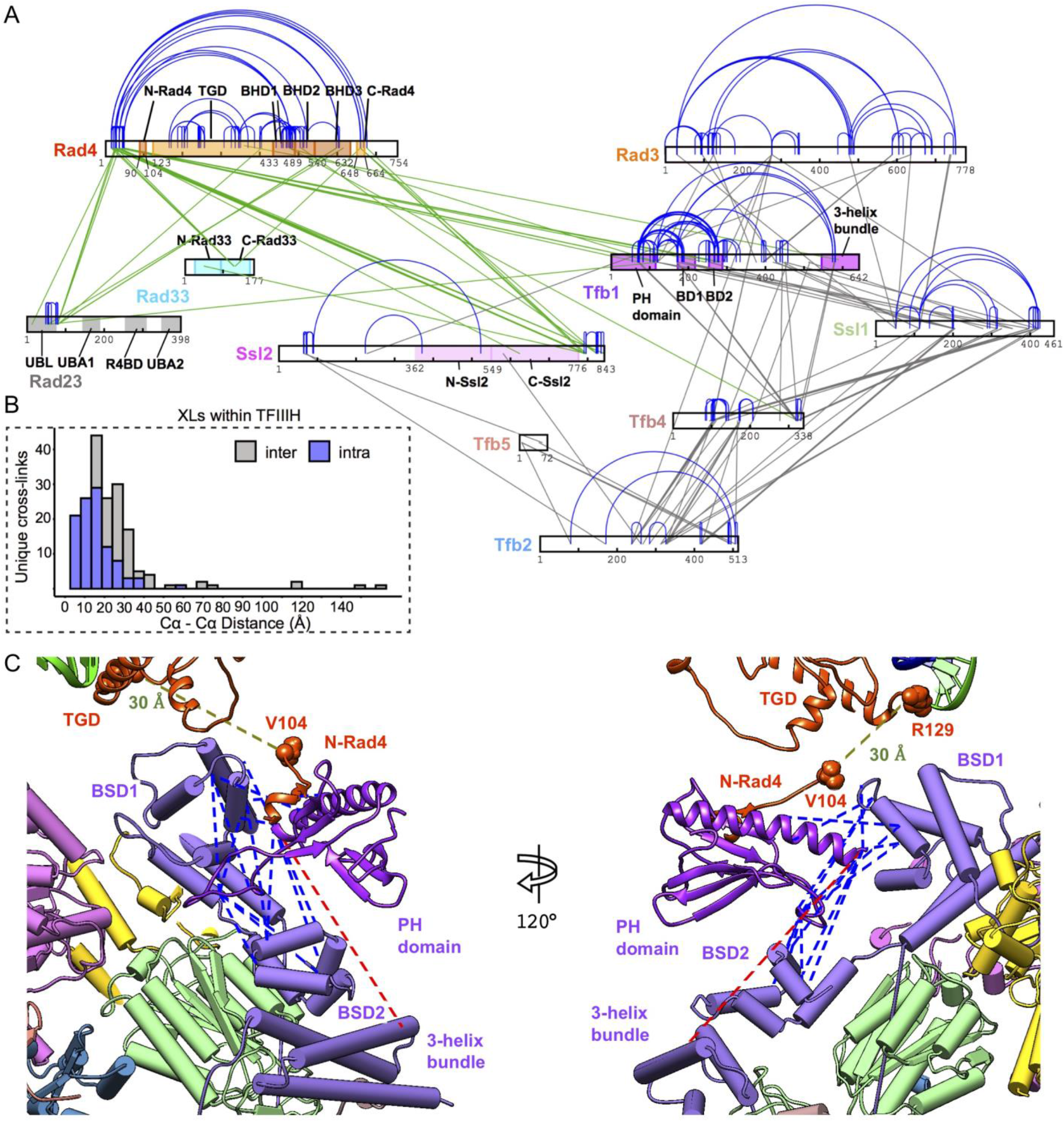
Cross-linking and mass spectrometry of TFIIH/Rad4-Rad23-Rad33/DNA. **(A)** Non-redundant cross-links identified for TFIIH/Rad4-Rad23-Rad33 on −24/+24 AAF-DNA as a network plot. Intra-subunit cross-links are shown in blue while inter-protein cross-links are shown in gray between TFIIH subunits and in green between Rad4-Rad23-Rad33 and TFIIH. Rad4, Rad23, Rad33 and TFIIH subunits that form cross-links with Rad4 are colored. **(B)** Cross-links were validated by mapping them on the near atomic structure of TFIIH in this study. 93% of cross-links were consistent with the 40 angstrom upper limit. **(C)** A tentative model of PH domain in TFIIH/Rad4-Rad23-Rad33/DNA complex. Of 16 unique cross-links to the PH domain that are mapped to Tfb1 domains, 15 cross-links are satisfied (blue) (Cα-Cα distances less than 40 angstroms), and 1 cross-link exceed 40 angstroms (red).

**Figure 3.**
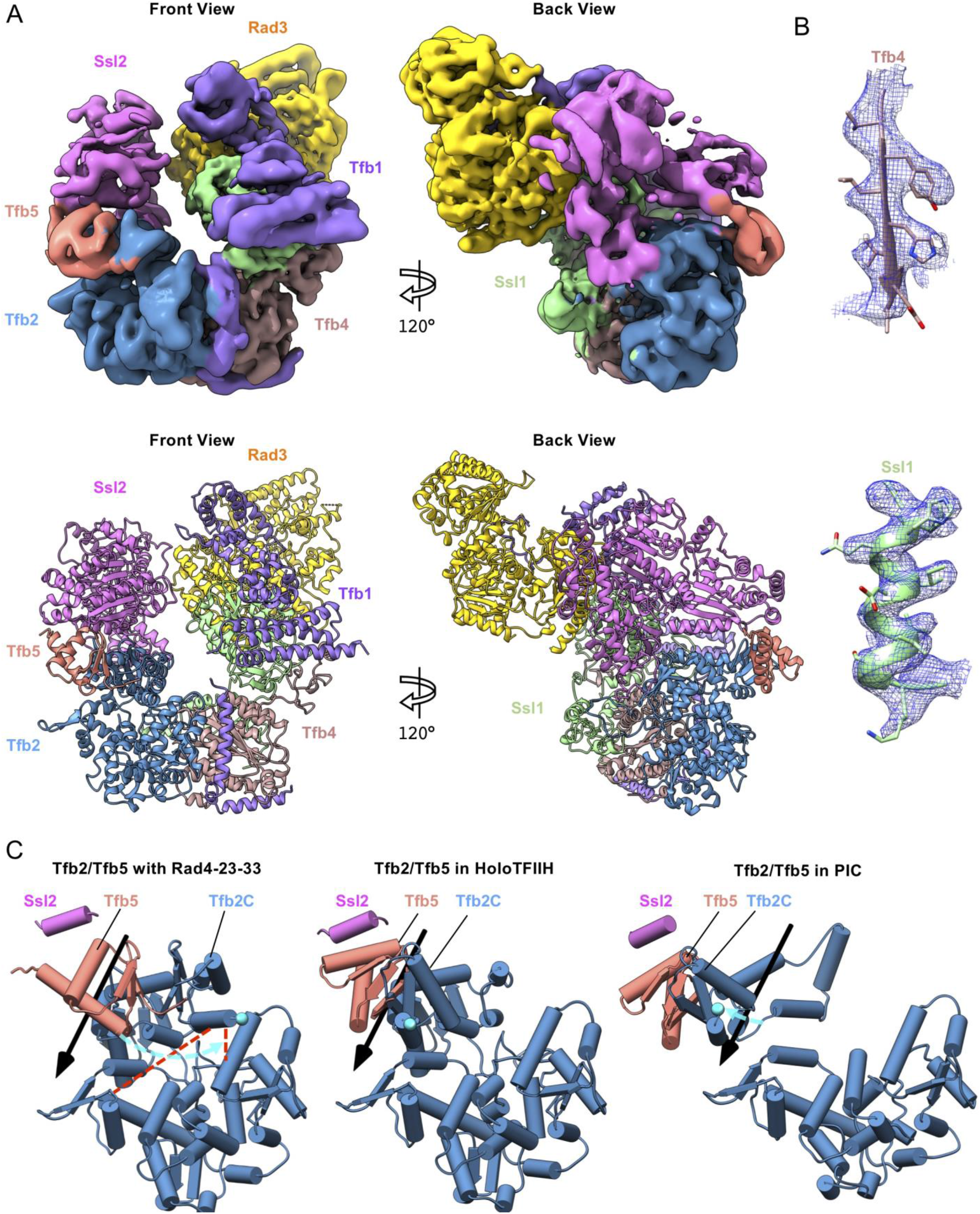
Structure of TFIIH in the TFIIH/Rad4-Rad23-Rad33/DNA complex. **(A)** Cryo-EM map (Map 1) and corresponding model of TFIIH in NER initiation complex shows clear density for each of the TFIIH subunits: Ssl2 (pink), Tfb5 (salmon), Tfb2 (blue), Tfb1 (gray), Tfb4 (brown), Ssl1 (green) and Rad3 (gold). Shown are front and 120-degree-rotated back views. **(B)** EM density map shows side-chain density for Ssl1 and Tfb4 subunits. **(C)** Conformational changes of Tfb2-Tfb5 upon binding Rad4-Rad23-Rad33 in NER. Relative to human holoTFIIH (PDB:6NMI, middle panel) and TFIIH in transcription PIC (PDB:5OQM, right panel), Tfb2C-Tfb5 undergoes a 60-degree rotation hinging at residues of 450-452 of Tfb2, accompanied by ~11 angstrom translation of Ssl2. R508 of Tfb2 is labeled (cyan spheres):

### Cryo-EM structure determination of TFIIH/Rad4-23-33/DNA complex

For cryo-EM studies, core TFIIH and Rad4-Rad23-Rad33 complexed with –24*/+24 AAF-DNA was prepared by glycerol gradient sedimentation in the presence of glutaraldehyde to enhance stability and minimize compositional heterogeneity ^53^. Aliquots of peak fractions embedded in ice disclosed fields of monodispersed particles in the presence of ATP as well as AMP-PNP (Figure S3A). We imaged ~1 million particles each with ATP or AMP-PNP using Titan Krios electron microscopes equipped with a K3 or K2 direct electron detector. 2D class averaging of particles in the presence of ATP yielded a set of homogeneous classes similar to one another except for differences in direction of view, some of which clearly showed the well-defined features of horseshoe-like TFIIH bound to the Rad4-Rad23-Rad33 complex (left panel of Figure S3B). In contrast, 2D class averages for the AMP-PNP-containing samples were more heterogeneous, hampering structural resolution, although some classes resembled TFIIH (right panel of Figure S3B). Noting the better homogeneity of the ATP-containing complexes, we selected ~250,000 images from this complex through 2D class averaging and subjected them to *ab initio* calculation of initial maps and subsequent 3D classifications. The resulting maps (Movie S1 and Table S1) showed clear division in two parts: a well-ordered TFIIH and a flanking disordered Rad4-23-33 (Figure S4). The most populated class resolved TFIIH at a nominal resolution of 3.86 Å, but with disordered density corresponding to the Rad4-Rad23-Rad33 complex bound to DNA (Map 1, EMDB-22587; Figures S4). To better resolve the disordered density, we refined a less populated class containing Rad4-Rad33/DNA at 9.2 Å resolution (Map 2, EMDB-22588; Figures S5), as well as a class containing Rad4 BHD3 (but not including BHD1-2 and TGD)-Rad33-DNA at 7.9 Å resolution (Map 3, EMDB-22576, Figure S5). Lastly multibody refinement ^54^ of Map 2 and Map 3 improved the local resolution of TFIIH to 8.1 and 7.0 Å, respectively, allowing us to better describe interactions between TFIIH and DNA (Figure S5). While the conformations of TFIIH in these three maps were similar to one another, they were significantly different from previous structures of TFIIH alone or engaged in transcription ^45,55^ (Figure 3C).

### Structure of TFIIH in the TFIIH/Rad4-Rad23-Rad33/DNA complex

Map 1 resolved TFIIH at a nominal resolution of 3.86 Å, the highest resolution for yeast TFIIH so far reported (Figures 3A-B). Local resolution calculations of Map 1 (Figure S3C) indicate that 6 of the 7 subunits of the core TFIIH (Rad3, Tfb1, Tfb2, Tfb4, Tfb5, and Ssl1) were determined at 3-6 Å resolution, while Ssl2 was determined at 6-8 Å resolution. Although ATP could not be resolved in the catalytic site of Ssl2, the structure of Ssl2 itself was a good match to the previous structure of the DNA-bound form of Ssl2 ^41–43^, characteristic of pre-translocation state of this family of helicases/translocases. Rad3 and its associated subunit Ssl1 were resolved at the highest resolution ~3-4 Å resolution within the complex. The DNA binding channel of Rad3 was occupied with the acidic plug of Tfb1 (residues 337-354) as previously observed in human XPD ^45^, indicating auto-inhibition of its DNA binding. A yeast homology model of TFIIH was generated using the human cryo-EM structure ^45^ as a template for further model building. While the overall subunit organization of TFIIH subunits was similar to that of human TFIIH solved by itself ^45^ (~4.6 Å RMSD), a significant difference was found in the position and orientation of Ssl2 and the domain that consists of Tfb5 (residues 2-72) and the C-terminal region of Tfb2 (Tfb2C; residues 412-513)^56^ (Figure 3C, Movie S2). In canonical TFIIH, the “open form” of Tfb5/Tfb2C was stabilized by the adjacent C-terminal helicase domain of Ssl2 (C-Ssl2, residues 549-776) through hydrophobic interactions between the N-terminal helix of Tfb5 (residues 14-26) and the hydrophobic helix (residues 563-571) of C-Ssl2 (light pink and brown red, respectively, Figure S3E). In the TFIIH/Rad4-23-33/DNA complex, Tfb5/Tfb2C were rotated ~60 °, and stabilized in the alternative position (“closed form”) through binding with the main body of TFIIH. There was little physical contact between Tfb5/Tfb2C and C-Ssl2 (salmon and pink, respectively, Figure S3E), and, due to the loss of this contact, the C-Ssl2 moved away by ~11 Å from the rest of TFIIH. Considering the primary role of Tfb5 in stimulating Ssl2 catalytic activity for the initial NER bubble formation in the presence of Rad4 (XPC) ^57^, the closed form of Tfb5/Tfb2C accompanied by repositioning of Ssl2 may represent an active form of TFIIH for initial NER bubble formation. TFIIH might undergo this conformational change induced by binding Rad4-Rad33/DNA (as described below). Indeed, an elongated density attributable to Rad4-Rad23-Rad33/DNA was seen on C-Ssl2, extending to Tfb1 across the complex, but was blurred due to the high mobility in Map 1. In contrast, the corresponding density was clearly seen in Map 2 as described below.

### Structure of Rad4-Rad23-Rad33 in the TFIIH/Rad4-Rad23-Rad33/DNA complex

In contrast to Map 1 that resolved TFIIH at high resolution but had a poor density for Rad4-Rad23-Rad33, Map 2 clearly showed the elongated density for the Rad4-Rad33 complex bound to DNA at 9.2 Å resolution (Figure 4A). While the TFIIH structure generally remained the same compared to that in Map 1, there was an additional density adjacent to BSD1 and BSD2 domains of Tfb1, attributed to the PH domain of Tfb1 (residues 1-114 of Tfb1) bound to an N-terminal segment of Rad4 preceding TGD (N-Rad4; residues 90–104 of Rad4)(purple in Figure 4A). The previous NMR model ^58^ was placed into the density, guided by with 9 and 6 cross-links formed with BSD1 and BSD2 of Tfb1, respectively (Figure 2C). Although the orientation of the PH domain remains tentative due to its limited resolution, the position evidently differed from that in transcription ^55^(Figure S2C). The crystal structure of Rad4–Rad23 bound to a 24 bp double-stranded DNA fragment containing a 6-4 photoproduct UV lesion (PDB:6CFI) ^23^ was a good match to the elongated density and was fitted as a rigid body (orange red in Figure 4A). In the resulting model, the TGD (residues 129-433 of Rad4) domain bound to DNA 3’ side of the lesion (mainly at positions +7 to +13) was in close proximity to the PH domain (left panel of Figure 4A) with the distance between the N-terminal end of the TGD (R129) and the C-terminal end of N-Rad4 (V104) being ~30 Å (Figure 2C). On the other hand, the BHD2-3 (residues 489-632 of Rad4) bound to a segment of DNA at the lesion (at positions –1 to +1) were located near Ssl2 (right panel of Figure 4A). When the DNA double helix 5’ side of the lesion ^23^ was extended with B-form DNA (Figure 5), a ~13 bp segment (at positions –15 to –3) was approximately in the position and orientation of DNA path along its binding groove between two ATPase domains of Ssl2 ^40–43^. The only minor adjustment needed for the modeling was a ~120 ° rotation (in a direction of unwinding) of the segment of DNA 5’ side of the lesion along its axis relative to the 3’ side (see next section).

**Figure 4.**
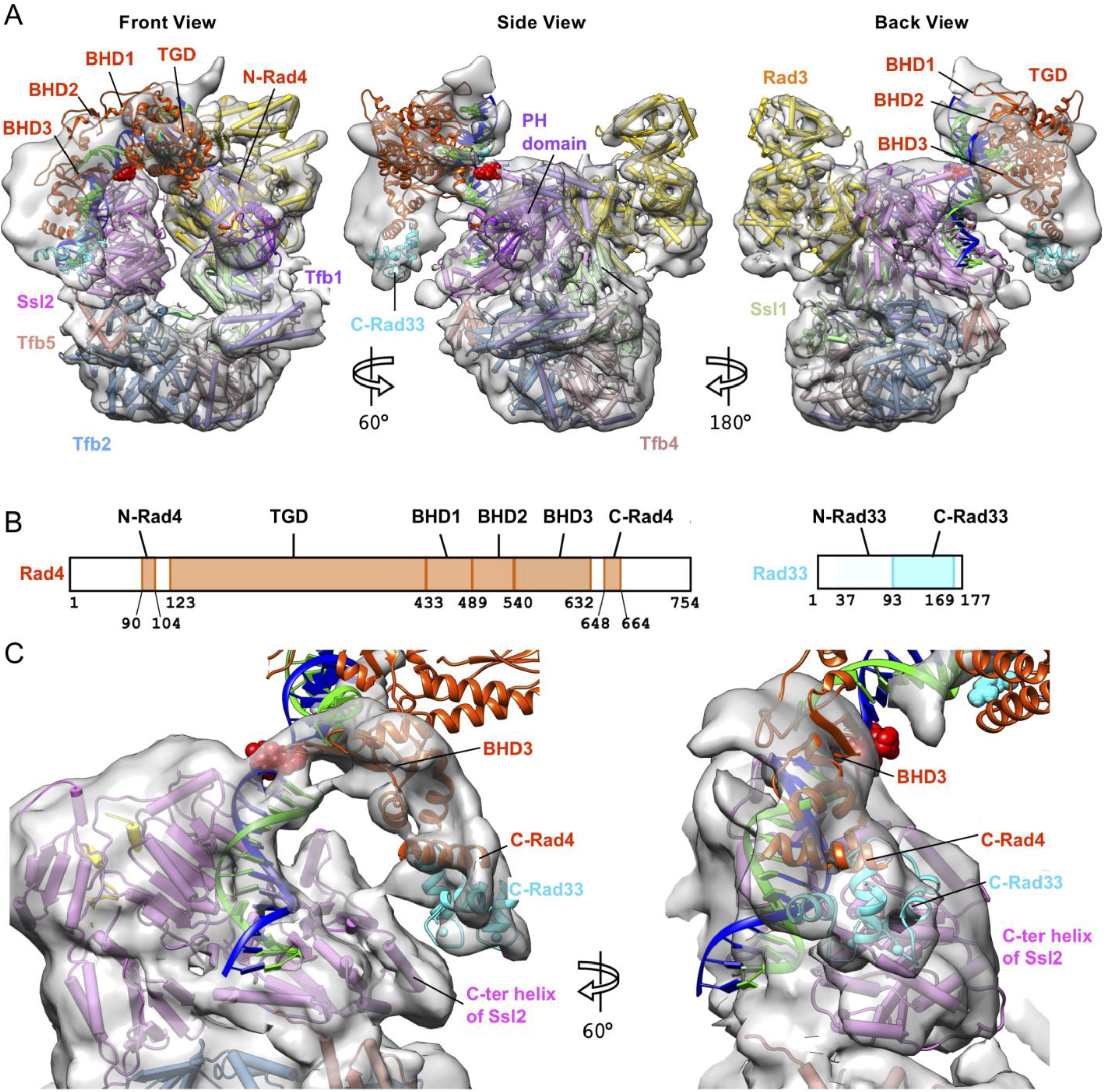
Structure of Rad4-Rad33 in the TFIIH/Rad4-Rad23-Rad33/DNA complex. **(A) Composite cryo-EM map of** Map 2 and Map 3 of TFIIH with Rad4-Rad23, seen in front (left), side (middle), and back (right) views. The crystal structure of Rad4-DNA complex (PDB:6CFI) was fitted into the density as a rigid body, with the lesion (6-4PP) in red sphere. Rad23 did not show corresponding density. **(B)** Schematic representation of domains of Rad4 and Rad33. Domains included in the model are colored. **(C)** 8.5 angstrom map (Map 3) refined by masking out TGD and BHD1-2. A homology model of Rad33 bound to C-Rad4 (PDB:2GGM) (cyan and orange red) was docked into a corresponding density, guided by cross-links of high confidence between C-Rad4 and the C-terminal region of Ssl2. The N-terminal half of Rad33 was not clearly seen. Seen in back view.

**Figure 5.**
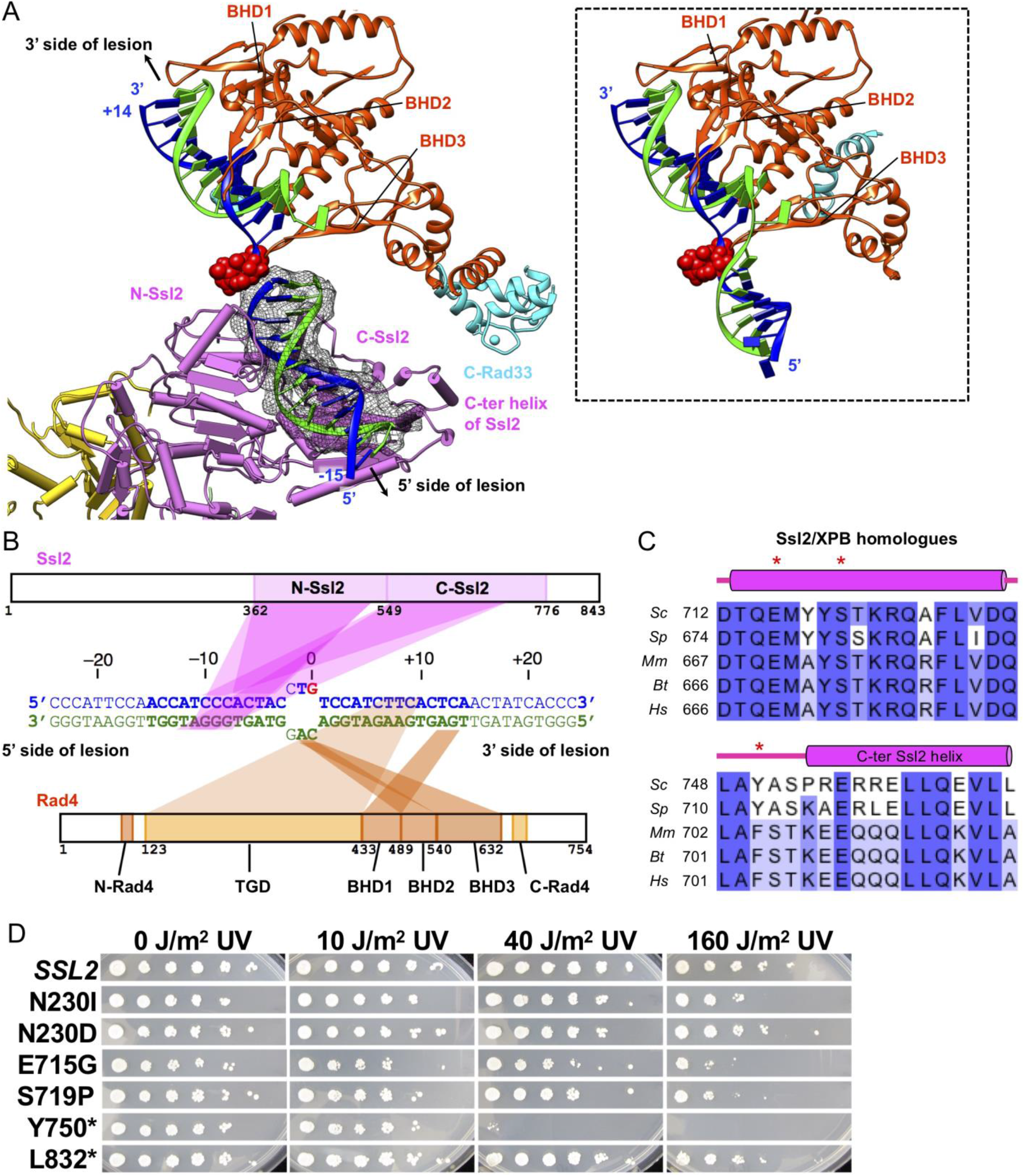
Path of damaged DNA in TFIIH-Rad4/23/33-DNA complex. **(A)** Zoomed view of protein-DNA interactions at the lesion (red sphere). ~13-bp DNA segment 5’ side of the lesion (black mesh) was identifiable on Ssl2 after multi-body refinement of Map 2. This segment of dsDNA is untwisted ~120 ° compared to that in Rad4-Rad23/DNA (PDB: 6CFI, inset), enabling simultaneous binding to Rad4 and Ssl2. **(B)** Schematic diagram of protein-DNA interactions suggested by the model. Nucleotide numbering follows the 5’-3’-direction on the damaged strand (blue) with respect to the AAF-dG at position +0. Nucleotides in the model are in bold. **(C)** Sequence alignment of C-Ssl2 and XPB with secondary structures between *Saccharomyces cerevisiae* (Sc), *Schizosaccharomyces pombe* (Sp), *Mus musculus* (Mm), *Bos taurus* (Bt), and *Homo sapiens* (Hs). Dots indicate positions for mutations of Ssl2 in (D). **(D)** Mutations of Ssl2 at the interface with Rad33 and Tfb5 confer UV sensitivity. Yeast cultures grown in YPD were diluted and plated. Uncovered plates were then irradiated with UV light (254 nm) at the indicated doses followed by incubation for 3 days. Asterisks indicate C-terminal deletion mutants at indicated positions.

Although the main body of Rad4 (residues 123-632) was in close proximity to the C-terminal helicase domain of Ssl2 (C-Ssl2, residues 549-776), there was apparently little direct contact with C-Ssl2. However, there was unassigned EM density bridging C-terminal end of Rad4 BHD3 (residue 632) and C-Ssl2 (Figure 4A), which indicated the presence of Rad33: Rad33 and human centrin 2 (CETN2) are EF-hand calcium-binding proteins and have been reported to interact with the conserved C-terminus region of Rad4 and XPC, respectively, and stimulate NER activity ^59–64^. To resolve this region, we further refined this map up to ~7.9 Å resolution (Map 3, Figures 4C and S5) using a mask that excludes the TGD and BHD1 domains of Rad4. This allowed us to reliably dock a homology model of the Rad33 C-terminal lobe (residues 94-169) bound to the Rad4 C-terminus region (C-Rad4; residues 648-664) into the corresponding density, guided by cross-links of high confidence between C-Rad4 and the C-terminal region of Ssl2 (*e.g.*, Rad4 S660 – Ssl2 K796) (Figure 2A) ^65,66^. The N-terminal lobe of Rad33 (residues 26-93) was not observed in the map. The conserved hydrophobic surface of C-Rad4 consisting of Thr651, Ile658, Leu662 ^66^ pack against hydrophobic residues of BHD3 (Phe585, Leu586, The624). Helix F (residues 127-136) of Rad33 was in close proximity to the C-terminal helix of C-Ssl2, which may contribute to the binding of the Rad4-23-33 complex to TFIIH. Consistent with our model, truncating the C-terminal 125 amino acids of human XPC (corresponding to residues ~617-754 of Rad4) impairs TFIIH binding *in vitro* ^52^. It should be noted that XPA (Rad14 in yeast) also binds this C-terminal helix of C-Ssl2 (Leu717 and Ala718) ^43^ (Figure S6), and thus may compete with XPC (Rad4) for binding Ssl2. Consistent with this, Rad14 did not significantly bind to or disrupt the TFIIH/Rad4-23-33/DNA complex (Figure S1F).

Altogether, our model shows that Rad4 binds TFIIH at two points of contact, N-Rad4 (residues 90–104) with the PH domain, and C-Rad4 (residues 648-664) with the C-terminal helix of C-Ssl2 through Rad33. The main body of Rad4 (residues 123-632 corresponding to TGD and BHD1-3) has no direct contact with TFIIH and is connected to N-Rad4 and C-Rad4 via short flexible linkers, which may account for the lower resolution of the EM map for Rad4 than for TFIIH.

### Untwisted DNA at the lesion for simultaneous binding to Rad4 and Ssl2

The localization of the lesion and the DNA path at its 3’ side were reliably deduced on the basis of fitting of the crystal structure of Rad4–Rad23 bound to a 24 bp dsDNA containing a 6-4 photoproduct UV lesion (PDB:6CFI) ^23^, whereas a ~13-bp segment of DNA double helix (at positions –15 to –3) 5’ side of the lesion was identifiable in the position and orientation of DNA path along its binding groove between two ATPase domains of Ssl2 ^40–43^ (Figure 5A). Compared to the DNA path in the crystal structure (PDB:6CFI), this segment of DNA 5’ side of the lesion was rotated by ~120 ° (in a direction of unwinding) along its axis relative to the 3’ side of the lesion (left vs right panels in Figure 5A). This untwisting required for simultaneous binding to Rad4 and Ssl2, is accommodated immediately 5’ side of the lesion at positions –2 and –3, where unwound strands were free of protein contacts. It should be noted that this segment of DNA double helix 5’ side of the lesion is highly mobile relative to the main body of Rad4-Rad23/DNA in the absence of TFIIH with a hinge point at position –2 ^22,23^, whereas the main body Rad4-Rad23-DNA shows little variability between crystal forms with different DNA substrates ^22,23^.

### Mutations of Ssl2 adjacent to the interface with Rad33 and Tfb5 confer UV sensitivity

To examine the structure-function relationship *in vivo*, a subset of novel *ssl2* alleles identified in screens for transcription-defective initiation factors were screened for UV sensitivities (Figures 5C-D). The mutant allele Y750*, a reconstruction of the mutant allele *SSL2-XP* ^67,68^ that eliminates 94 C-terminal amino acid residues (residues 750-843), confers extreme UV sensitivity, in good agreement with localization of the binding sites for Rad33 and Rad14 in this region (Figures 5C-D). Two additional alleles, E715G and S719P, which are located at the interface with the C-terminal loop of Tfb5 (Figure S5E), have moderate UV sensitivity. Other ssl2 mutants were either not or were much less sensitive to UV light, suggesting separation of NER and transcription related functions, or a much greater sensitivity of transcription-related phenotypes than of UV-sensitivity. Overall, UV sensitive *ssl2* mutations were localized on binding sites for NER factors (Rad14, Rad33, and Tfb5).

## Discussion

Structural and mechanistic studies of NER involving TFIIH have been hampered by difficulties in preparing the relevant NER factors and their assemblies on DNA in sufficient quality and quantity *in vitro*. With highly purified NER factors, we investigated the assembly of the initiating NER complex and developed a procedure for isolating homogeneous complexes containing the 7-subunit core TFIIH and Rad4-Rad23-Rad33. Here, we present the first structural model of TFIIH recruited by Rad4(XPC) to damaged DNA (AAF-DNA), as determined by the 3.9–9.2 Å resolution cryo-EM maps and XL-MS analysis, which mutually reinforce conclusions from each. In this structure, ~30-bp stretch of damaged DNA duplex could be visualized as it bound to Rad4 on one end and the Ssl2 DNA translocase subunit of TFIIH on the other. This model is in good agreement with a prevailing model derived from biochemical studies with regard to Rad4(XPC)-TFIIH interactions and positioning of DNA lesion ^17,48,51,52,69,70^. Importantly, it provides key insights into how NER initiates as TFIIH is recruited to damaged DNA by Rad4(XPC).

Biochemical studies such as DNA footprinting assays have long been indicated that NER involves progressive DNA unwinding (‘opening’) at the lesion as NER factors are recruited to the site in an orderly fashion ^47,48^. For various NER lesions, the initial ‘opening’ is first carried out during lesion recognition as Rad4 bends and unwinds DNA around the lesion and flips out two damage-containing nucleotide pairs from the DNA duplex ^22,23^. However, how the next level of ‘opening’ and lesion verification involving TFIIH may be executed remained unclear.

The DNA in the present TFIIH/Rad4-23-33/DNA structure shows that the unwinding is extended by at least one base pair (positions from 0 to –2) compared to that in Rad4-Rad23/DNA (positions from 0 to –1). The additional unwinding is a consequence of the simultaneous binding of DNA with Rad4 and Ssl2 (Figure 5). Achieving such simultaneous binding may require ATP hydrolysis: samples assembled without ATP but with AMP-PNP suffered from structural heterogeneity, which could be due to failing to achieve stable dual binding (Figure S3B)^17^. The relatively small additional unwinding depicted in our structure indicates that the structure captures an early step of TFIIH engagement in NER and that DNA unwinding is needed even for the initial loading of TFIIH.

Moreover, the organization of molecules together with their known biochemical functions provides insights that extend beyond this early stage. In particular, our structure sheds light on how the DNA ‘opening’ can be further extended in the downstream process involving the translocase activity of Ssl2 (Figure 6C). In RNA polymerase II transcription, Ssl2 is bound to downstream DNA in the transcription preinitiation complex. Using a molecular wrench mechanism, Ssl2 translocates along the DNA that induces DNA melting ~20 bp downstream of the TATA box. This in turn allows the delivery of template strand to the active center of the polymerase for the transition from a closed to open promoter complex ^41,71–74^. In the TFIIH/Rad4-23-33/DNA structure, Ssl2 is bound at the 5’ side of the lesion, poised to extend the DNA bubble around the lesion in a manner analogous to how it forms a transcription bubble (Figure 6C): the NER bubble would extend as Ssl2 reels downstream (5’-side) dsDNA towards Rad4, while Rad4 remains stably bound with upstream (3’-side) DNA at the lesion site (covering positions –1 to +13, Figure 5B). Consistent with this model, previous permanganate footprinting assays exhibited a KMnO_4_ hyperreactive region around positions 0 to –10 on the damaged strand upon full engagement of XPB ^48^. Along with Ssl2, Rad4 is likely to play an important role in NER bubble formation. As previously noted, position and orientation of Rad4 with respect to TFIIH is held in place by its dual interaction with Ssl2 and Tfb1 of TFIIH through its flexible C- and N-termini, respectively. Without Rad4 holding the 3’ side of the DNA as an anchor, the DNA may freely rotate around its helical axis to release the torsional stress created by Ssl2 translocation, instead of directing it to induce DNA unwinding.

**Figure 6.**
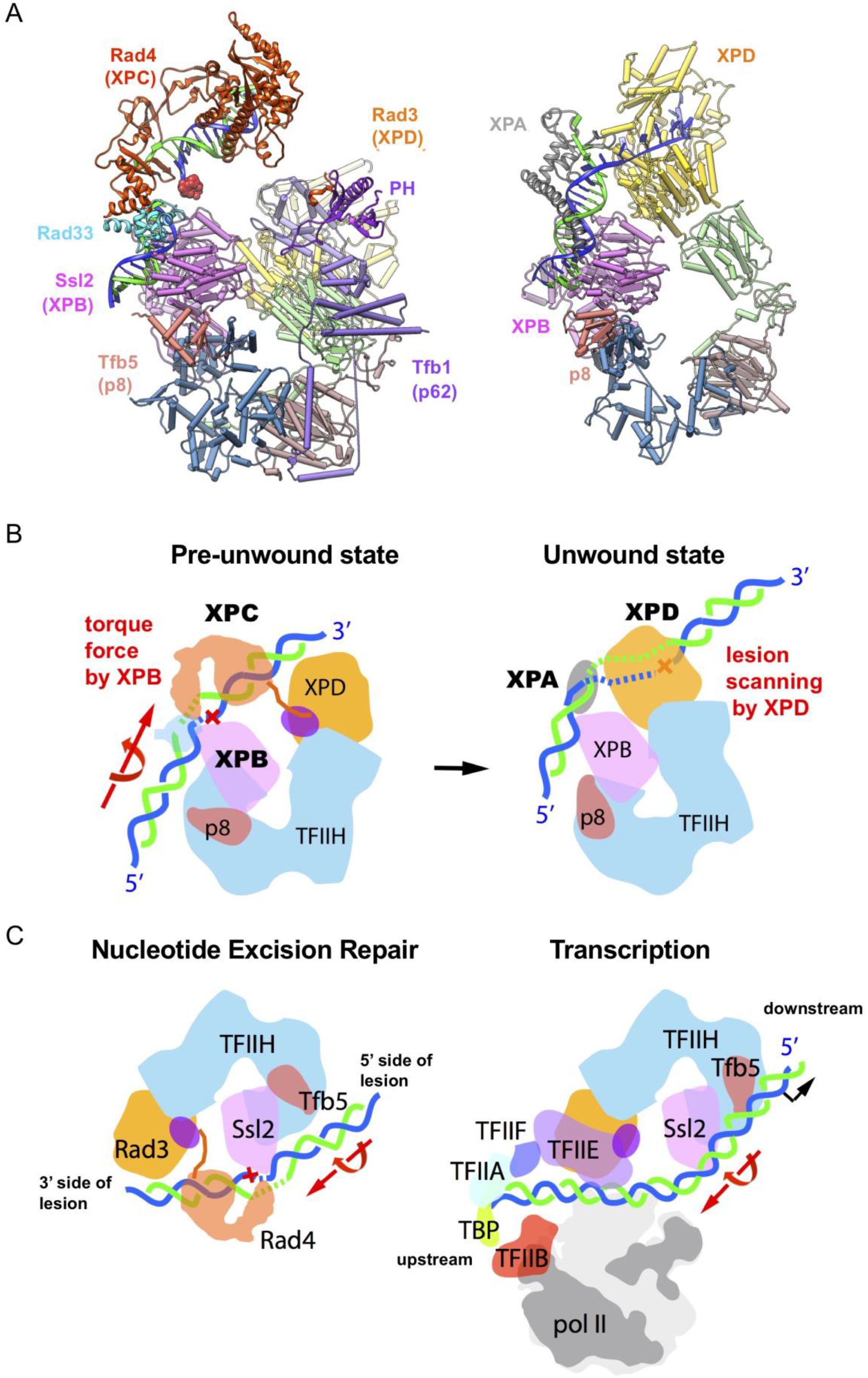
Two structures of TFIIH and schematic model of unwinding of DNA for NER. **(A)** Ribbon representation of the yeast pre-unwound state (left) and previous structure of the human (artificially) unwound state (right, PDB:6RO4). Two structures are aligned by superimposing Ssl2 and XPB. **(B)** A model for unwinding damaged DNA by TFIIH, XPC, and XPA in NER. Lesion bound XPC is stabilized on TFIIH primarily through DNA binding to XPB (left). XPC is then positioned near XPD for lesion scanning and verification by a concerted movement of XPD. The torque force by the translocase activity of XPB opens a bubble near damage, facilitating delivery of the damage DNA to XPD in an unwound form suitable for binding in the DNA binding cleft (right). This unwinding step is also key to the lesion verification step in which a bulky lesion stalls and blocks the progression of XPD helicase unwinding, thereby allowing the incision complex to be stably formed. **(C)** A comparison between NER initiation (left) and pol II transcription initiation (right). Two forms of TFIIH in NER initiation and transcription initiation are aligned. DNA in blue and green is rotated and translocated by Ssl2 in an ATP-dependent manner, in the directions indicated by the large red arrows.

We also posit that an enlarged NER bubble will then create a single-stranded DNA platform sufficient to engage with Rad3 and downstream factors such as Rad14(XPA) for lesion verification. XPD is supposed to scan the damaged strand by its 5’-3’ helicase activity in order to verify a bona fide bulky NER lesion and prevent futile excision of non-bulky, undamaged DNA. In the current model, however, Rad3 was not in contact with DNA, indicating that further conformational changes must take place to deliver the damaged strand to Rad3. Recently, a cryo-EM structure mimicking the Rad3-engaged “unwound” state was reported, where core TFIIH and XPA was assembled on a fork DNA substrate containing unduplexed, single-stranded DNA segment that permitted TFIIH and XPA loading while bypassing the requirement for XPC (Figures 6A and S6) ^43^. Compared with canonical TFIIH structures, XPD(Rad3) in this structure moved ~80 Å to bind to the ssDNA segment in its DNA-binding channel, whereas XPB kept the same position while bound to a dsDNA segment. When our yeast structure (“pre-unwound” state) was aligned with this human structure (“unwound” state) by superposing Ssl2 on XPB, the ~80 Å movement of XPD(Rad3) from the pre-unwound to unwound states did not cause major steric clash with Rad4. The repositioned XPD was also suitably positioned to bind to the damaged DNA strand in its DNA binding cleft (with 5′ to 3′ ssDNA binding polarity from left to right in Figures 6A-B). This suggests that the structural transition from pre-unwound to unwound state may happen while retaining Rad4 (Movie S3).

In contrast, the same superposition showed that Rad33 and XPA(Rad14) would encounter steric clash with each other as both bind to the C-terminal helix of C-Ssl2 (Figure S6). This suggests that Rad33 and Rad14 are mutually exclusive when they bind to Ssl2, assuming Rad14-Ssl2 interactions are conserved with XPA-XPB interactions. Biochemical studies have suggested that Rad33 and Rad14 functions at distinct steps in NER: the former in promoting DNA damage recognition with XPC(Rad4) and early TFIIH loading ^59,60^ and the latter in activating XPD(Rad3) helicase after initial TFIIH loading by XPC(Rad4) ^17,28,43^. The transition between the pre-unwound and unwound stages of the NER initiation may thus entail the binding of Rad14 to TFIIH-DNA while displacing Rad33 from Ssl2.

Together, our study sets the stage for more detailed mechanistic questions for subsequent steps. For instance, how does the translocase activity of XPB and helicase activity of XPD in TFIIH orchestrate DNA unwinding and subsequent lesion scanning for damage verification ^17,28^? How does XPA stimulate this process ^28,75^? As XPD is ~70 Å away from DNA at the pre-unwound state, how is the unwound damaged strand delivered to the DNA binding cleft of XPD? Previous data have pointed to the release of CAK (TFIIK) as TFIIH engages with DNA damage to function in NER ^38,39^. The TFIIH-XPA-DNA structure by Kokic et al., also shows that TFIIK binding is incompatible in this stage where DNA is engaged with XPD and is already substantially unwound. Our structure, on the other hand, indicates that in the initial stage of TFIIH recruitment, TFIIK may still be bound. Therefore, the dissociation/association of TFIIK must be orchestrated at a later event before or as XPA joins, and the exact mechanism remains unknown. We speculate that this event is also likely key to switching TFIIH’s role between transcription and NER. Additionally, our structural study has significant clinical implications. Many XP/CS/TTD mutations of TFIIH ^76^, can be mapped at the interface with NER factors. This will open new doors to understand various NER-linked diseases and to develop novel therapeutic interventions against them.

## Methods

### Protein purification

The Rad4–Rad23 and complexes were prepared as previously described ^22^. Briefly, the Hi5 insect cells co-expressing the Rad4–Rad23 complex were harvested 2 days after infection. After lysis, the proteins were purified using His-Select Nickel agarose resin (Sigma) and anion exchange chromatography (Source Q, GE Healthcare), followed by thrombin digestion and cation exchange (Source S, GE Healthcare) and gel-filtration (Superdex200, GE Healthcare) chromatography. The final sample was concentrated by ultrafiltration to ∼13 mg/ml in 5 mM bis–tris propane–HCl (BTP-HCl), 800 mM sodium chloride, 5 mM dithiothreitol (DTT), pH 6.8.

Holo-TFIIH were purified from yeast as previously described ^29^ with minor modifications. In short, tfb6Δ strain containing TAP tags on TFIIH subunits Tfb4 and Ssl2 was grown by fermenter (Eppendorf) in 100 L of YPAD medium to OD 10.0. Whole cell lysate was prepared by bead beating in Buffer A (50 mM HEPES pH 7.6, 1 mM EDTA, 5% glycerol, 400 mM potassium acetate, 2-mercaptoethanol, and protease inhibitors). Following the addition of 100 mM ammonium sulfate and 0.12% PEI, lysed cells were stirred for 1 h and centrifuged, and then the cleared lysate was loaded onto an IgG column. The column was washed with 5–10 column volumes of buffer 300 (50 mM HEPES pH 7.6, 1 mM EDTA, 5% glycerol, 300 mM potassium acetate, 2mM DTT, and protease inhibitors) then resuspended in buffer 300 and allowed to settle overnight. IgG beads were washed by batch with another 10 column volumes of buffer 300 over the course of 24 hours. TFIIH was treated with TEV for 15 hours in buffer 300, eluted from the IgG column, spin concentrated then loaded onto an UnoQ column (Bio-Rad). TFIIH was eluted by salt gradient of concentration from 300 mM to 1.2 M potassium acetate. Fractions containing 10 component holo-TFIIH, six component coreTFIIH and five component TFIIH lacking Ssl2 s were concentrated separately, aliquoted and flash frozen.

Rad33 was overexpressed in bacteria and purified by the following method. Rosetta(DE3) (Fisher Scientific) were transformed with pETDuet-RAD33 plasmid and grow overnight. 2L of Terrific Broth (TB) was inoculated with overnight culture and grown to OD600=1.0. Cells were induced with 1mM Isopropyl β-D-1-thiogalactopyranoside (IPTG) for 4 hours and harvested by centrifugation (Sorvall LYNX 6000, ThermoFisher Scientific) at 8,000 rpm for 10 minutes. Cells were resuspended in lysis buffer (50 mM HEPES (pH 7.6), 500 mM sodium chloride, 5 mM 2-mercaptaethanol, 20 mM imidazole, 5% (v/v) glycerol and protease inhibitors) and lysed by sonication. Lysate was clarified by centrifugation at 40,000 rpm for 45 minutes and loaded onto a HisTrap column (GE Healthcare) equilibrated in nickel buffer A (50 mM HEPES (pH 7.6), 500 mM sodium chloride, 5 mM 2-mercaptaethanol, 20 mM imidazole, 5% (v/v) glycerol). Column was washed with 10 column volumes of 3% nickel buffer B (50 mM HEPES (pH 7.6), 500 mM sodium chloride, 5 mM 2-mercaptaethanol, 750 mM imidazole, 5% (v/v) glycerol) and then eluted with a linear gradient of nickel buffer A to nickel buffer B over 90 mL. Peak fractions were pooled, diluted ~5 fold with buffer 0 (20 mM HEPES, 2 mM DTT) and loaded onto a Q column (GE Healthcare). Rad33 was then eluted from the Q column using a linear gradient of Q buffer A (20 mM HEPES, 100 mM sodium chloride, 2mM DTT, 5% (v/v) glycerol) to Q buffer B (20 mM HEPES, 1M sodium chloride, 2mM DTT, 5% (v/v) glycerol). Peak fractions were pooled, buffer exchanged to buffer 300 (20 mM HEPES, 300 mM potassium acetate, 5 mM DTT) during concentration and frozen.

Rad2 was overexpressed in bacteria and purified by the following method. Rosetta(DE3) (Fisher Scientific) were transformed pGVD52-RAD2HisSumo. Transformed cells were grown in TB media at 37°C, induced with 0.4 mM IPTG at OD600 0.9 and grown overnight at 16°C. Cells were harvested by centrifugation (Sorvall LYNX 6000, ThermoFisher Scientific) at 8,000x rpm for 10 minutes and resuspended in lysis buffer that contained 50 mM Tris-HCl (pH 7.5), 500 mM sodium chloride, 40 mM imidazole, 5% (v/v) glycerol and 5 mM 2-mercaptoethanol and protease inhibitors. Cells were lysed by sonication and lysate was clarified by centrifugation at 40,000 rpm for 45 minutes. The supernatant was loaded on a HisTrap column (GE Healthcare) equilibrated in lysis buffer. The column was then washed with lysis buffer that contained 1.5 M sodium chloride and then buffer A (50 mM Tris-HCl (pH 7.5), 500 mM sodium chloride, 40 mM imidazole, 5% (v/v) glycerol, 5 mM 2-mercaptoethanol) and then eluted from the column with a linear gradient of gradient of nickel buffer A to nickel buffer B over 90 mLs. 6xHis-Ulp1 protease was added to the protein-containing fractions, which were then dialyzed overnight in a buffer that contained 20 mM Tris-HCl, 250 mM sodium chloride, 5% (v/v) glycerol, 5 mM 2-mercaptoethanol. Dialyzed samples were re-applied on a HisTrap column, and protein that contained flow-through was loaded onto a Heparin HP column (GE Healthcare). Rad2 was eluted from by a linear gradient of heparin buffer A (20 mM Tris-HCl, 250 mM potassium acetate, 5% (v/v) glycerol and 2 mM DTT) to heparin buffer B (20 mM Tris-HCl 1M potassium acetate, 5% (v/v) glycerol and 2 mM DTT). Peak fractions were pooled and concentrated.

Rad14 was overexpressed in bacteria and purified by the following method. Rosetta(DE3) (Fisher Scientific) were transformed pET-RAD14His. Transformed cells were grown in TB media supplemented with 10 μM zinc chloride at 37°C, induced with 1 mM IPTG at OD600 1.0 and grown for three hours at 25°C. Cells were harvested by centrifugation (Sorvall LYNX 6000, ThermoFisher Scientific) at 8,000x rpm for 10 minutes and resuspended in lysis buffer that contained 100 mM Tris-HCl (pH 8.0), 500 mM sodium chloride, 20 mM imidazole, 10% (v/v) glycerol and 2mM DTT, 10 μM zinc chloride and protease inhibitors. Cells were lysed by sonication and lysate was clarified by centrifugation at 40,000 rpm for 45 minutes. The supernatant was loaded on a HisTrap column (GE Healthcare) equilibrated in nickel buffer A (100 mM Tris-HCl (pH 7.5), 100 mM sodium chloride, 20mM imidazole, 5% (v/v) glycerol) and eluted from the column with a linear gradient of gradient of nickel buffer A to nickel buffer B (buffer A + 500 mM imidazole) over 90 mLs. Protein containing fractions were then loaded onto a Q HP column (GE Healthcare) and eluted from by a linear gradient of Q buffer A (100 mM Tris (pH 7.5), 100 mM sodium chloride, 5% (v/v) glycerol and 5 mM DTT) to Q buffer B (100 mM Tris (pH 7.5), 100 mM sodium chloride, 5% (v/v) glycerol and 5 mM DTT). Peak fractions were dialyzed into buffer 300 (20 mM HEPES (pH 7.6), 200 mM potassium acetate, 5% glycerol, 1 mM DTT), concentrated and frozen.

### N-AAF DNA Synthesis and Purification

N-acetoxyacetomino fluorinated DNA aducts were purified via a modified protocol from previous described ^16^. 1.25 nmole of oligonucleotides with single G residues were incubated with 0.1 mM N-acetoxy-2-acetylaminofluorene in TE buffer at 37°C for 3 h in a light sensitive container. The top AAF-modified oligonucleotides were separated from unmodified ones by C18-reversed phase high-performance liquid chromatography (Beckman Ultrasphere 5 μm; φ4.6 × 250 mm). The column was equilibrated sample loaded with a mixture of 100 mM triethylamine acetate (Sigma) and acetonitrile (95:5). Bound oligonucleotides were eluted by linearly increasing the acetonitrile concentration from 10% up to 30%. Complementary bottom unmodified oligonucleotides were purified by Urea-PAGE. Top and bottom oligos were mixed, annealed and then the double stranded damaged or undamged product was purified by size exclusion chromatography (Superose 6, GE Healthcare) in buffer 300 (20 mM HEPES pH 7.6, 300 mM potassium acetate, 5 mM DTT).

### Electromobility Shift Assays

Purified DNA substrates were labeled by mixing 15 pmoles of AAF DNA, with T4 PNK (New England BioLabs), PNK buffer and P^32^ gamma-ATP for 90 minutes at 37°C. Excess P^32^ gamma-ATP was then separated from labeled AAF DNA by G50 spin column (GE Healthcare). Labeled AAF DNA was mixed 1:4 with unlabeled AAF DNA prior to use for electromobility shift assays (EMSA).

EMSAs were performed by mixing equal concentrations (250 nM) of Rad4-Rad23, coreTFIIH, and labeled/unlabeled AAF DNA in buffer 100 (20 mM HEPES, 100 mM potassium acetate, 2 mM magnesium acetate, 5 mM DTT) with additional factors as indicated. Reactions were incubated at room temperature for 20 minutes. Reactions were run on 0.2x TBE agarose gels at 4°C for 2 hours. Gels were vacuum dried (BioRad) and imaged by phosphorimager.

### Rad2/Rad14 Competition Assays

To prepare samples for competition assays, 120 pmoles of Rad4-Rad23, 120 pmoles of Rad33 and 120 pmoles of −24/+24 AAF were mixed and dialyzed to assembly buffer (20 mM HEPES pH 7.6, 300 mM potassium acetate, 2 mM magnesium acetate, 5% (v/v) glycerol, 5 mM DTT) for 4 hours. Dialyzed Rad4-Rad23-Rad33/AAF was then added to 0.83 nmoles of coreTFIIH with 5 mM ATP or 2 mM AMP-PNP 2 mM magnesium acetate and 120 pmole of Rad14 or Rad2. Reactions were incubated for 60 minutes. TFIIH/Rad4-Rad23-Rad33/AAF +/− Rad14/Rad2 were then sedimented at 52,000 rpm for 6 hours and 35 minutes in a 10 to 40% glycerol gradient (20 mM HEPES pH 7.6, 150 mM potassium acetate, 2 mM magnesium acetate, 4 mM DTT, 2 mM ATP or 1 mM AMP-PNP) at 4°C using a Beckman SW 60 Ti rotor. Glycerol gradients were prepared using a Gradient Master device (BioComp Instruments). Samples were then fractionated with a Piston Gradient Fractionator (BioComp Instruments). Protein distribution was analyzed by TCA precipitation of fractions followed by SDS-PAGE analysis. Gel images were scanned and then densitometry analysis was performed in ImageJ ^77^ and data were plotted in Prism (GraphPad).

### Cryo-EM Sample Preparation and Data Collection

To prepare samples for cryo-EM analysis, 1.15 nmoles of Rad4-Rad23, 1.18 nmoles of Rad33 and 1.15 nmoles of −24*/+24 AAF were mixed and dialyzed to assembly buffer (20 mM HEPES, 300 mM potassium acetate, 2 mM magnesium acetate, 5% (v/v) glycerol, 5mM DTT) for 4 hours. Dialyzed Rad4-Rad23-Rad33/AAF was then added to 0.83 nmoles of coreTFIIH, with 5 mM ATP and 2mM magnesium acetate and incubated for 60 minutes. TFIIH/Rad4-Rad23-Rad33/AAF complex was then prepared as previously described ^53^: samples were sedimented at 52,000 rpm for 6 hours and 35 minutes in a 10 to 40% glycerol gradient (20 mM HEPES, 150 mM potassium acetate, 2 mM magnesium acetate, 4 mM DTT, 2 mM ATP) at 4°C using a Beckman SW 60 Ti rotor. Glycerol gradients were prepared using a Gradient Master device (BioComp Instruments). When cross-linking was used for EM analysis, 0.125% (v/v) glutaraldehyde was added to the 40% glycerol solution before gradient preparation. Samples were then fractionated with a Piston Gradient Fractionator (BioComp Instruments). Cross-linking reactions were quenched by the addition of glycine-HCl buffer (pH 7.5) to a final concentration of 40 mM. Peak was analyzed by TCA precipitation of uncrosslinked fractions followed by SDS-PAGE analysis. Peak crosslinked fractions were concentrated and then frozen. A similar procedure was followed for preparation of AMP-PNP state of TFIIH/Rad4-Rad23-Rad33/AAF complex by replacing ATP with AMP-PNP in all of the steps.

To prepare cryo-EM grids, purified TFIIH/Rad4-Rad23-Rad33/AAF was thawed and dialyzed in EM buffer (20 mM HEPES pH 7.6, 150 mM potassium acetate, 5 mM DTT) for 30 minutes immediately prior to making grids. 2uL of dialyzed sample was then applied to glow-discharged (1 min, easiGlow, Pelco) R2/2 200- or 300-mesh Quantifoil holey carbon grids (Electron Microscopy Sciences). The grids were subsequently blotted for 2 or 3 seconds, respectively, using Whatman Grade 41 filter paper (Sigma-Aldrich) and flash-frozen in liquid ethane with a Leica EM CPC manual plunger (Leica Microsystems). EM grids were prepared in batches and the freezing conditions were optimized by screening on a FEI TF20 microscope operating at 200 kV and equipped with a FEI Falcon III direct electron detection camera at the Electron Microscopy Research Lab (University of Pennsylvania).

Cryo-EM specimens were imaged at two different settings. Dataset 1 was collected at University of Massachusetts Medical School was collected using a FEI Titan Krios G3i transmission electron microscope operating at 300 kV, equipped with a K3 direct electron detector (Gatan) and a Bioquantum energy quantum filter (Gatan). Data was collected by SerialEM ^78^ by image shift and at a nominal magnification of 105,000x in super-resolution mode (pixel size of 0.415 Å) at a defocus range between 0.8 and 2.8 μm. A total of 6234 images were over the course of one and a half days. Exposures were 2.6 seconds with movies divided into 30 frames at a targeted nominal dose of 50 e^−^/Å2.

Dataset 2 was collected at the NIH National Center CryoEM Access and Training (NCCAT) at the New York Structural Biology Center (NYSBC) using a FEI Titan Krios transmission electron microscope operating at 300 kV, equipped with a K2 Summit direct electron detector (Gatan) and a Bioquantum energy quantum filter (Gatan). Data was collected by Appion ^79^ by image shift and at a nominal magnification of 130,00x in super-resolution mode (pixel size of 0.504 Å) at a defocus range between 0.8 and 2.8 μm. A total of 2073 images were over the course of two days. The exposure time was 5 s at a nominal dose of 42 e^−^/Å2.

### Image processing and 3D reconstruction

A combination of software including cryoSPARC v2.12.4 ^80^, Relion 3.0.8 ^81^, and Relion 3.1 were used to process both cryo-EM datasets. Datasets were processed independently as follows. The K3 dataset (Umass) dataset was motion-corrected with MotionCorr2 ^82^ then imported into cryoSPARC for CTF correction with CTFFIND4 ^83^. A total of 1,159,898 particles were picked by blob picking, and two rounds of reference-free 2D classification were performed to remove particles that lacked clear features (Supplementary Figure 3B), resulting in a subset of 250,223 particles. This subset was then subjected to initial model calculation followed by four rounds of heterogenous refinement then non-uniform refinement all in cryoSPARC. The resulting map (7.1 angstrom resolution) showed clear TFIIH density with noticeable rotation of Tfb2C-Tfb5. This map was low pass filtered and then used as an initial model for 2 rounds of heterogeneous refinement of the entire 1,159,898 particle dataset which resulted in four maps with reasonable TFIIH features. The best map, containing 196,883 particles, yielded a 4.9 angstrom resolution map after non-uniform refinement in cryoSPARC. These particles were then transferred to Relion 3.0.8 for an additional round of 3D classification. The best class contained 56,101 particles and yielded a 4.8 angstrom resolution map after 3D auto-refinement in Relion 3.0.8. After multiple rounds of CTF refinement and Bayesian polishing a 3.86 angstrom resolution map was obtained (Map 1).

A similar method as described above was used for processing the K2 (NCCAT dataset). The K2 dataset (Umass) dataset was motion-corrected with MotionCorr2 ^82^ then imported into cryoSPARC for CTF correction with CTFFIND4 ^83^. A total of 595,544 particles were picked by blob picking, and a single round of reference-free 2D classification were performed to remove particles that lacked clear features, resulting in a subset of 155,529 particles (subset 1). These particles were imported into Relion 3.1 and subjected to 3D classification without mask (Supplementary Figure S4B). One of the class contained 63,672 particles with clear density for both Rad4 and TFIIH. Auto refinement of this map using a soft mask containing entire Rad4 and TFIIH yielded a map of 9.25 angstrom resolution (Map 2). To obtain higher resolution of Rad4 BHD3 and TFIIH, another subset of 73,146 particles (subset 2) from cryoSPARC was imported into Relion 3.1 and subjected to 3D classification with a soft-edge mask containing Rad4 BHD3 and TFIIH. Auto refinement of the best resolved class containing 32,858 particles using the same soft-edge mask yielded a map of 7.95 angstrom resolution (Map 3).

### Model building and refinement

To build the model of TFIIH in the high-resolution map (Map 1), we started with rigid body fitting of TFIIH subunits from the EM structure of yeast TFIIH (PDB:5OQJ) ^55^ using UCSF Chimera ^84^. To model large portions of the Ssl2 N-terminus and Tfb2, which are absent from that model but visualized in our map, the human TFIIH structure (PDB: 6NMI) ^45^ was used as a template for homology modeling with MODELLER ^85^. Tfb5-Tfb2C (residues 438-508) were well outside the density and fitted into the density as a rigid body in UCSF Chimera. The resulting model for seven yeast TFIIH subunits was then used as a template for structure refinement using RosettaCM ^86^ and manual refinement with Coot (Emsley and Cowtan, 2004). The final model statistics are found in Supplemental Table 1.

For the TFIIH/Rad4-Rad23-Rad33/AAF model, the TFIIH model obtained from Map 1 was docked into the map containing density for Rad4-Rad33/AAF (Map 2).Then a crystallographic model of a Rad4-Rad23 bound to a 6-4 photoproduct UV lesion (PDB: 6CFI) was docked as a rigid body into the EM map (Map 2) without any deviations. Rad23 was not included in the model due to the lack of the corresponding density. DNA 5’ side of the lesion was aligned to DNA path on XPB (PDB: 6RO4). A homology model of Rad33 bound to C-Rad4 was generated using the crystal structure of human Centrin 2 bound to XPC peptide (PDB: 2GGM) as a template ^65^, and was fitted into the corresponding density as a rigid body ^84^. PH domain bound to N-Rad4 (PDB:2M14) was placed in the corresponding density and refined as a rigid body with Phenix1.16 ^87^. The final (only most plausible) model was chosen based on the fitting score. Figures were made using a combination of UCSF Chimera ^84^ and UCSF ChimeraX ^88XXXX^.

### Cross-linking mass spectrometry sample preparation

150 μg of TFIIH/Rad4-Rad23-Rad33/AAF was prepared by the same method for EM analysis. Assembled TFIIH/Rad4-Rad23-Rad33/AAF was diluted to a concentration of 1 mg/mL with buffer 300 (20 mM HEPES (pH=7.6), 300 mM potassium acetate, 5% glycerol, 5 mM DTT) and then mixed with 50 mM (final 6 mM) disuccinimidyl dibutyric urea (DSBU) (Thermo Fisher Scientific). Crosslinking reaction was incubated on ice for 2 hours and then was quenched by adding 1M (final 50 mM) of ammonium bicarbonate, and then further stopped by TCA precipitation. Crosslinked proteins were precipitated with 20% (w/v) trichloroacetic acid (TCA, Sigma) on ice for 90 minutes. Proteins were pelleted by centrifugation at 21,000 x g for 15 min and washed with 10% TCA in 100mM Tris-HCl and then with acetone (Fisher Scientific). The solvent was decanted, the pellet air-dried and then stored at −80°C for analysis by mass spectrometry.

Crosslinked proteins were resuspended in 50 μl of resuspension buffer (2.5% SDS and 50 mM triethylammonium bicarbonate (TEAB) final concentrations) and reduced with final 10 mM DTT (US Biological) for 30 min at 30 °C, followed by alkylation with final 50 mM iodoacetamide (Sigma Aldrich) for 30 min at 30 °C. The proteins were captured in an S-Trap™ mini column (Protifi, C02-mini) to remove contaminants, salts, and detergents and concentrate the proteins in the column for the efficient digestion then digested with trypsin (Thermo Fisher Scientific) in 1:10 (w/w) enzyme/protein ratio for 1 h at 47 °C. Peptides eluted from this column, with 50 mM TEAB, 0.2% formic acid, and 60% acetonitrile in order, were vacuum-dried and resuspended with the peptide fractionation buffer [(70% (v/v) LC-MS grade water (Thermo Fisher Scientific), 30% (v/v) acetonitrile (ACN, Thermo Fisher Scientific) and 0.1 % (v/v) trifluoroacetic acid (TFA, Thermo Fisher Scientific)]. To enrich for crosslinked peptides, peptides were first fractionated using an AKTA Pure 25 with a Superdex 30 Increase 3.2/300 column (GE Life Science). System was flowed at a rate of 30 μL/min of fractionation buffer and 100 μL fractions were collected from the beginning of the sample-loop injection to 1.5 column volumes. Based on the elution profile, fractions containing enriched crosslinked peptides of higher molecular masses were vacuum-dried and resuspended in 0.1% (v/v) TFA in LC-MS grade water for mass spectrometry analysis.

Fractions were analyzed by a Q-Exactive HF mass spectrometer (Thermo Fisher Scientific) coupled to a Dionex Ultimate 3000 UHPLC system (Thermo Fischer Scientific) equipped with an in-house made 15 cm long fused silica capillary column (75 μm ID), packed with reversed phase Repro-Sil Pure C18-AQ 2.4 μm resin (Dr. Maisch GmbH, Ammerbuch, Germany). Elution was performed by the following method: a linear gradient from 5% to 45% buffer B (90 min), followed by 90% buffer B (5 min), and re-equilibration from 90% to 5% buffer B (5 min) with a flow rate of 400 nL/min (buffer A: 0.1% formic acid in water; buffer B: 80% acetonitrile with 0.1% formic acid). Data were acquired in data-dependent MS/MS mode. Full scan MS settings were as follows: mass range 300−1800 m/z, resolution 120,000; MS1 AGC target 1E6; MS1 Maximum IT 200. MS/MS settings were: resolution 30,000; AGC target 2E5; MS2 Maximum IT 300 ms; fragmentation was enforced by higher-energy collisional dissociation with stepped collision energy of 25, 27, 30; loop count top 12; isolation window 1.5; fixed first mass 130; MS2 Minimum AGC target 800; charge exclusion: unassigned, 1, 2, 3, 8 and >8; peptide match off; exclude isotope on; dynamic exclusion 45 sec. Raw files were converted to mgf format with TurboRawToMGF 2.0.8 ^89^.

### Crosslinked peptide search

To identify and validate crosslinked peptides the search engine MeroX 2.0.1.4 ^90^ was used. MeroX was run in RISEUP mode, with default crosslinker mass and fragmentation parameters for DSBU and the following adjusted parameters: precursor mass range, 1,000–10,000 Da; minimum precursor charge 4; precursor and fragment ion precisions 5.0 and 10.0 ppm, respectively; maximum number of missed cleavages 3; carbamidomethylation of cysteine and oxidation of methionine, as fixed and variable modifications, respectively. Results were filtered for score (>10) and false discovery rate, FDR (<1%). The in-house R script was utilized to automate the execution of MeroX to submit to a cluster, combine the results, prioritize K-K pair if lysine miscleavage presents and localization score ties, and filter the results to show only unique pairs with the best scores. The Xlink Analyzer plug-in ^91^ for UCSF Chimera was used for visualization of the crosslinks and placement of the PH domain of Tfb1.

### *ssl2* phenotypic analysis

A subset of novel *ssl2* alleles identified in screens for transcription-defective initiation factors were screened for UV sensitivity. These alleles were identified based on randomized mutagenesis of *SSL2* and screening for phenotypes we have found to correlate with initiation defects, sensitivity to mycophenolic acid and activation of a transcriptional fusion of *HIS3* and the *IMD2* promoter. These phenotypes are previously described ^92^. For UV sensitivity testing, liquid cultures of relevant strains were grown overnight in YPD. 10-fold serial dilution series of each strain were spotted onto minimal medium lacking leucine (SC-Leu) and exposed to indicated doses of 265 nm UV light using an FB-UVXL-1000 UV crosslinker (Fisher). Plates were incubated at 30°C in the dark for three days and then imaged. Yeast media were as described previouslyb^95^ with modifications previously described ^93^. Sequence alignments were generated by clustal omega multi sequence alignment and colored by sequence conservation in Jalview ^94^.

## Acknowledgments

We thank the University of Massachusetts CryoEM Core Facility, Drs. Chen Xu, KangKang Song and Kyounghwan Lee for their constant support and assistance in data collection. We thank Dr. Sudheer Molugu and Electron Microscopy Resource Laboratory at the University of Pennsylvania for use of equipment and assistance with cryo-EM sample screening. Some of this work was performed at the National Center for CryoEM Access and Training (NCCAT) and the Simons Electron Microscopy Center located at the New York Structural Biology Center, supported by the NIH Common Fund Transformative High Resolution Cryo-Electron Microscopy program (U24 GM129539), and by grants from the Simons Foundation (SF349247) and NY State. We thank Drs. Yi-Wei Chang and Shrawan Mageswaran for assistance with data analysis by cryo-electron tomography. We would like to thank Dr. Geoffrey Dann for his assistance in purifying AAF substrates. This research was supported by NIH R01-GM123233 to K.M., NIH R01GM120450 to C.D.K., NSF grant MCB-1412692 and NIH grant R21-ES028384 to J.-H.M, NIH grant P01CA196539 to B.A.G.; NIH training grants T32-GM008275 to T.V.E and T32-GM071339 to H.J.K. Computational resources were supported by NIH Project Grant S10OD023592.

## Author contributions

T.V.E., J-H.M., K.M. designed the experiments. T.V.E. and Y.S. prepared samples. T.V.E. prepared cryo-EM samples and analyzed the data. T.V.E., J-H.M., and K.M. built a model. H.J.K. and B.A.G. performed and analyzed XL-MS. T.Z., S.B., and C.K. performed in vivo analysis. T.V.E, J-H.M., C.K., and K.M. wrote the paper and prepared the figures for publication.

## Competing interests

Authors declare no competing interests.

## Data and materials availability

Cryo-EM maps and models were deposited in the Electron Microscopy Data Bank (EMDB-XXXX for Map 1, EMDB-XXXXX for Map 2, EMDB-XXXXX for Map 3). The atomic coordinates were deposited in the Protein Data Bank (accession code: XXXX, XXXX). Crosslinking Mass-Spectrometry data of TFIIH/Rad4-Rad23-Rad33/AAF complex was deposited in XXXXXXXX.

---

**Supplemental Figure 1.**
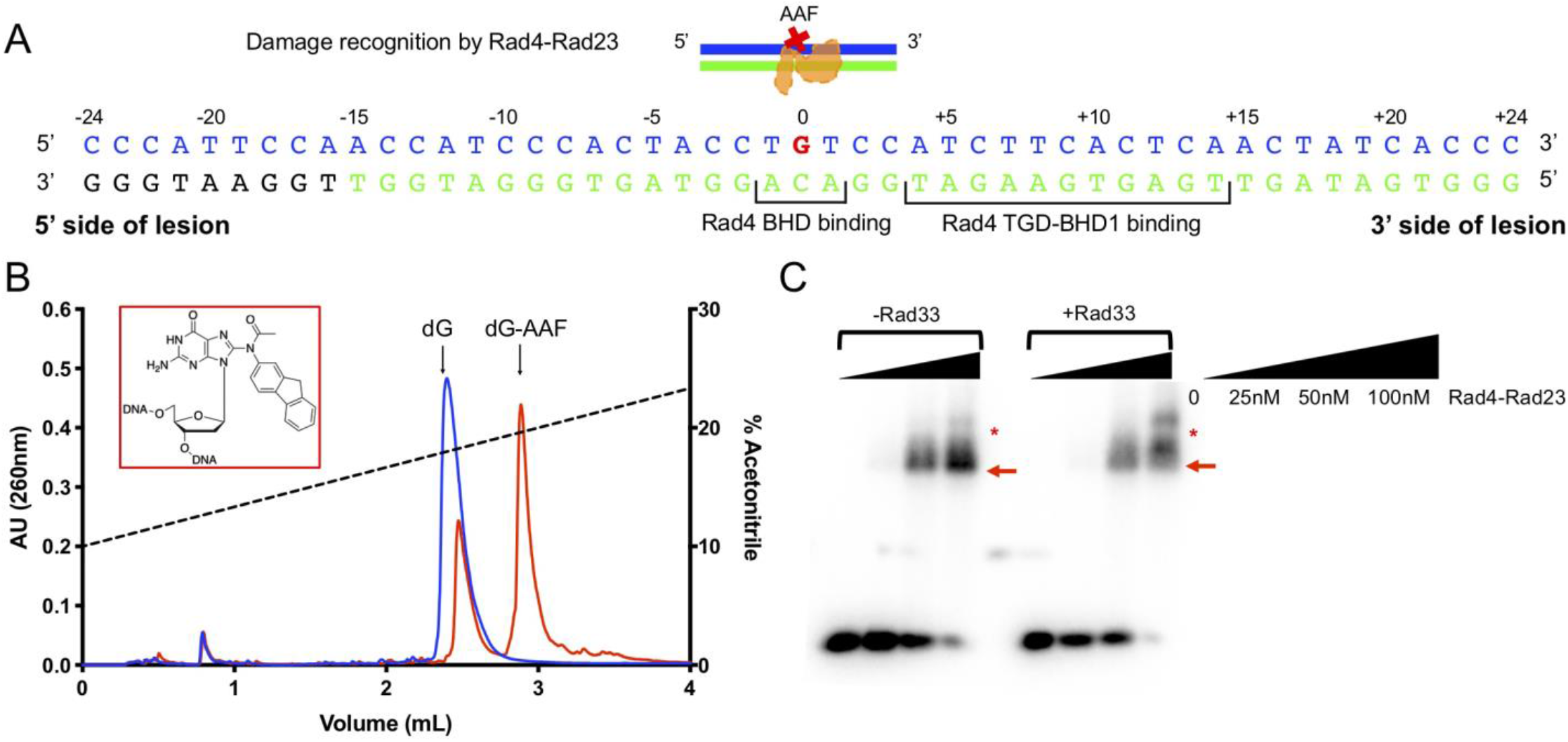
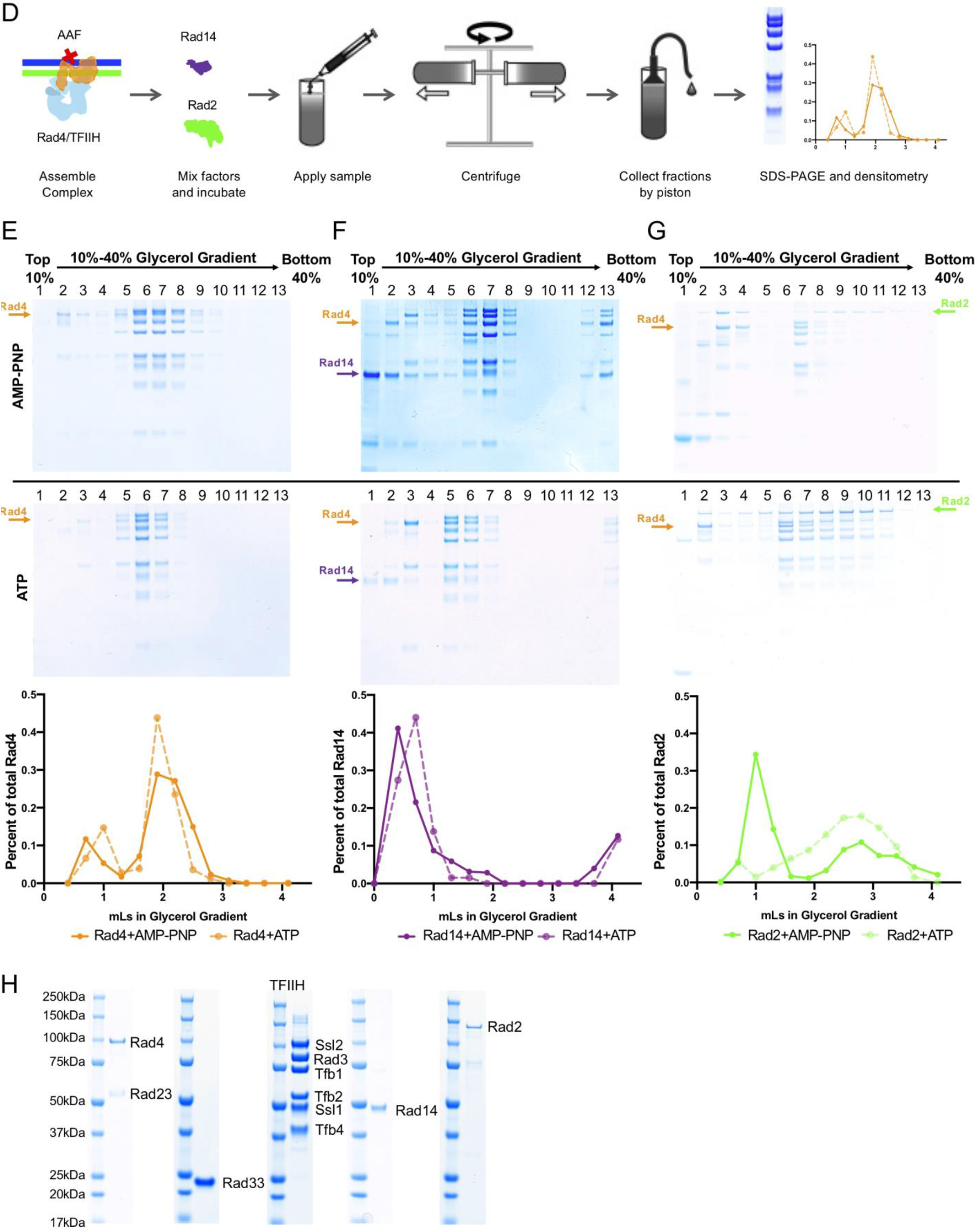
**(A)** Schematic illustration of Rad4 bound to a lesion (AAF)-containing DNA duplex (upper panel) and the sequence of the −24/+24 AAF-DNA used as a DNA template in this study (lower panel). AAF-dG is in red at position 0. **(B)** Separation of AAF-dG oligonucleotide (red) from single stranded dG oligo (blue) by HPLC reverse phase chromatography. N-Acetoxy-2-acetamidofluorene reacted specifically at guanosine (inset) under conditions described in Methods. **(C)** EMSA of TFIIH with increasing amounts (0-100 nM) of Rad4-Rad23 with −24/+24 DNA in the presence or absence of 100 nM of Rad33. Stoichiometrically bound complexes are indicated by red arrows, whereas some non-specific/non-stoichiometric complexes are indicated by red asterisks. **(D)** Schematic illustration of competitive binding assay with Rad14 and Rad2. **(E)** Isolation of the TFIIH/Rad4-Rad23-Rad33 complex with −24*/+24 AAF in the presence of AMP-PNP (top) or ATP (middle). **(F-G)** The TFIIH/Rad4-23-33/DNA complex was combined by addition of 6-fold molar excess of Rad14 (F) or Rad2 (G) and then subjected to glycerol gradient sedimentation in the presence of AMP-PNP (top) or ATP (middle). Proteins were analyzed by SDS-PAGE analysis. Proteins of interest (Rad4, Rad14 or Rad2) were quantified by densitometry (bottom). TFIIH-Rad4/23/33-DNA complex was stable during sedimentation in the presence of AMP-PNP or ATP in fractions 6-8 (E). Rad14 failed to bind TFIIH/Rad4-23-33/DNA complex in either AMP-PNP or ATP (F). Rad2 did not show robust binding with AMP-PNP (G, top) but did bind with ATP, and facilitated dissociation of Rad4-Rad23 (G, middle). **(H)** SDS-PAGE analysis of purified NER proteins.

**Supplemental Figure 2.**
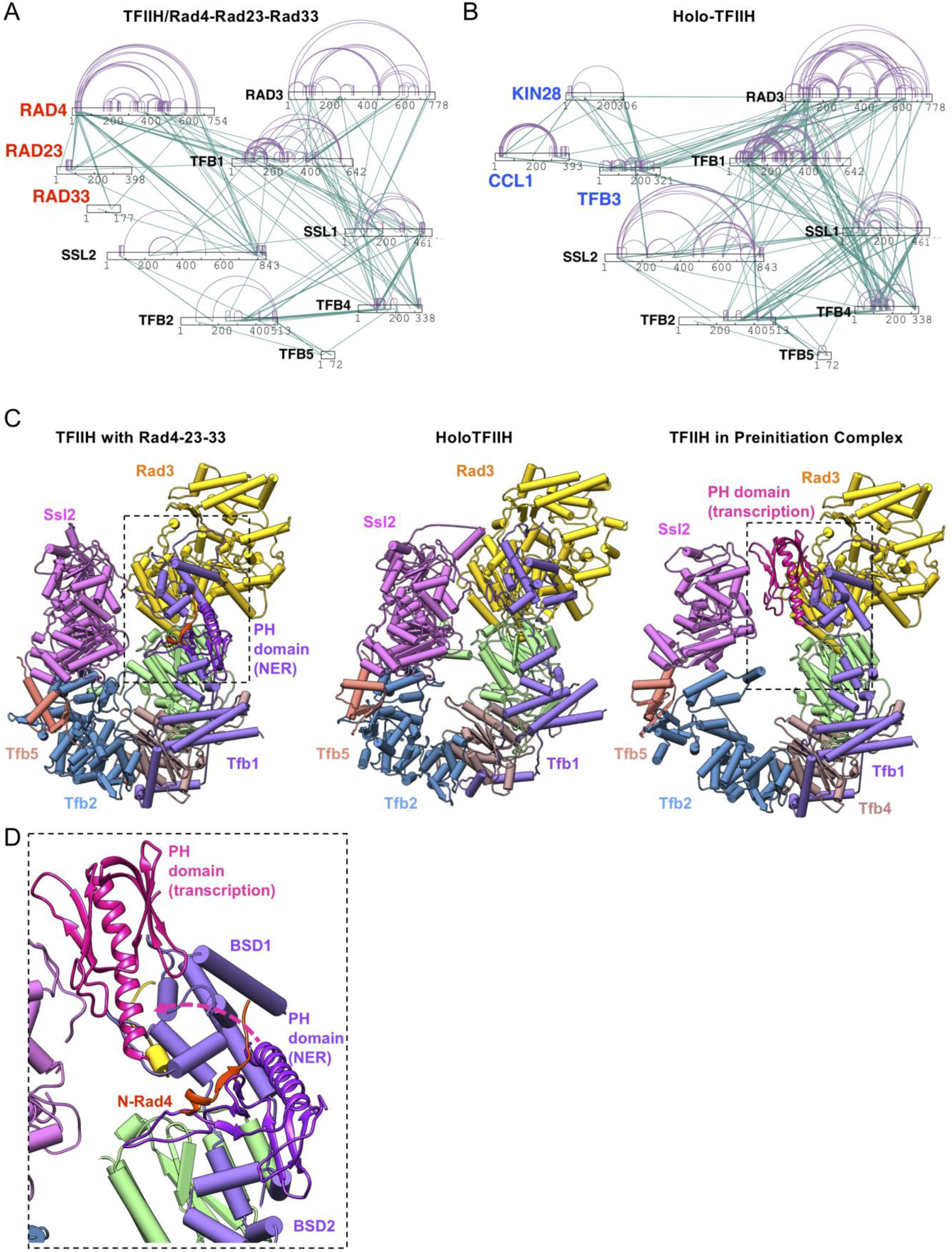
**(A-B)** Cross-links identified for TFIIH/Rad4-23-33/DNA (A) and holo-TFIIH containing TFIIK (B) as a network plot. Intra-subunit cross-links are shown in purple while inter-protein cross-links are shown in green. **(C)** Comparison of structures of TFIIH between with Rad4/23/33 (left), HoloTFIIH (middle, PDB:6NMI) and in the preinitiation complex (right, PDB:5OQM). **(D)** Comparison of Tfb1 PH domain positioning in transcription initiation (magenta, PDB:5OQJ) and NER initiation (purple). Structures aligned by superposition of the Tfb1 BSD and the 3-helix bundle domains.

**Supplemental Figure 3.**
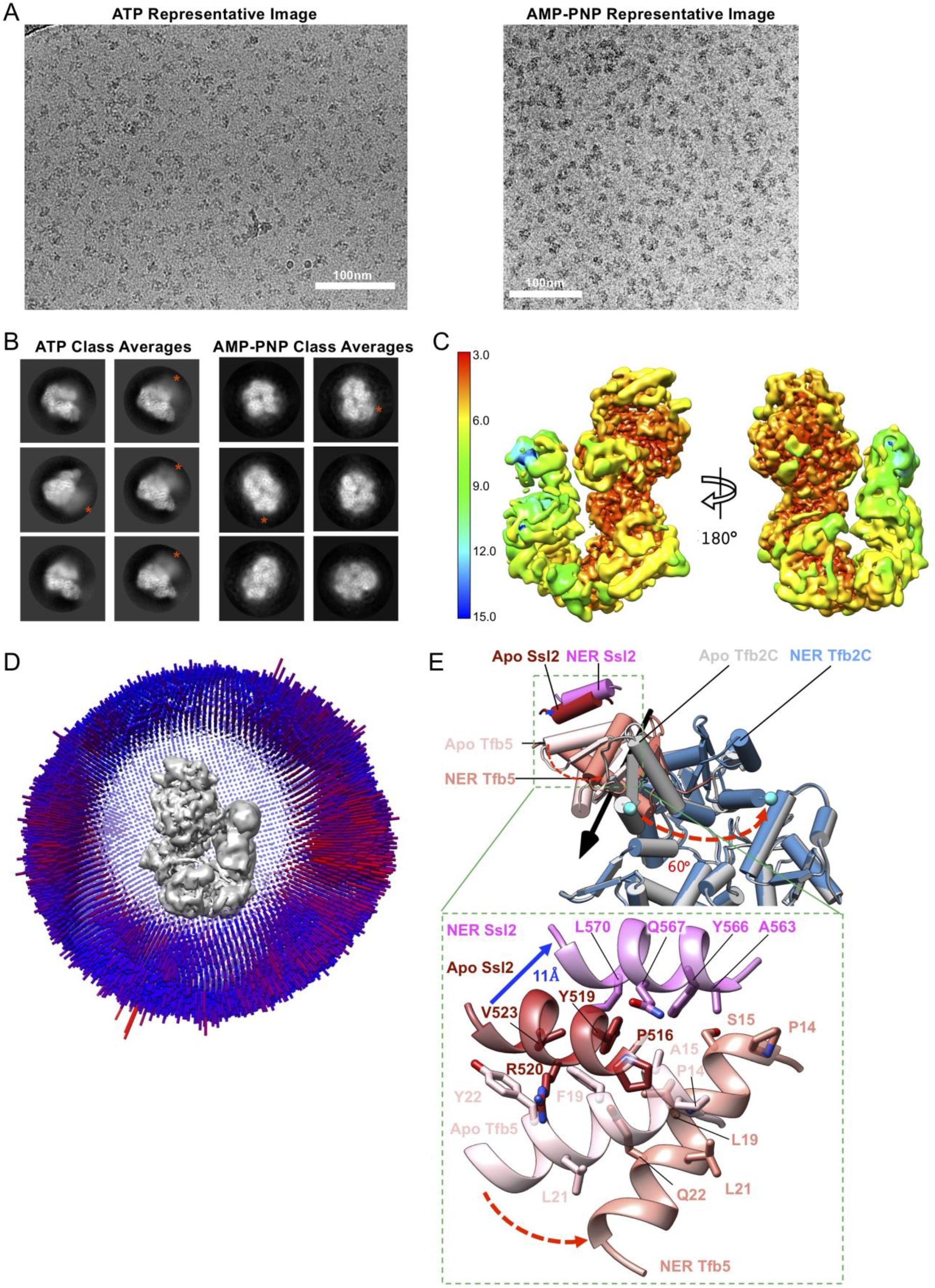
**(A)** Representative electron micrograph of TFIIH/Rad4-23-33/DNA complex assembled in the presence of AMP-PNP or ATP. Scale bar represents 100nm. **(B)** Representative 2D class averages obtained from reference-free 2D classification of TFIIH/Rad4-23-33/DNA complex by Relion 3.0. Rad4 (red asterisk) in the samples assembled in the presence of ATP (left) or AMP-PNP (right) shows alternative binding locations relative to TFIIH. **(C)** Local resolution of the core TFIIH within the ternary complex in the ATP state as calculated by Blocres at FSC=0.5. Ssl1, Tfb4 and Rad3 represent a high-resolution core of the complex with local resolution ~3-4 angstroms. Ssl2 and Tfb2C-Tfb5 formed a structural module variable in position and orientation relative the rest of TFIIH with local resolution ~7-9 angstroms. **(D)** Euler plot distribution of particle views in the final reconstruction, where the view representation increases from blue to red. **(E)** Conformational changes of Tfb2-Tfb5 upon binding Rad4-Rad23-Rad33. Relative to human TFIIH (PDB:6NMI), Tfb2C-Tfb5 undergoes a 60-degree rotation hinging at residues of 450-452 of Tfb2, accompanied by ~11 angstrom translation of Ssl2. R508 of Tfb2 is labeled in cyan spheres. Enlarged view of interaction between the N-terminal helix of the NER specific subunit Tfb5 (salmon) and the hydrophobic helix of C-Ssl2 (pink). Structure of human core TFIIH (PDB:6NMI) was aligned by superimposing Tfb2 and p52. p8 and XPB of human core TFIIH are shown in light pink and brown red, respectively.

**Supplemental Figure 4.**
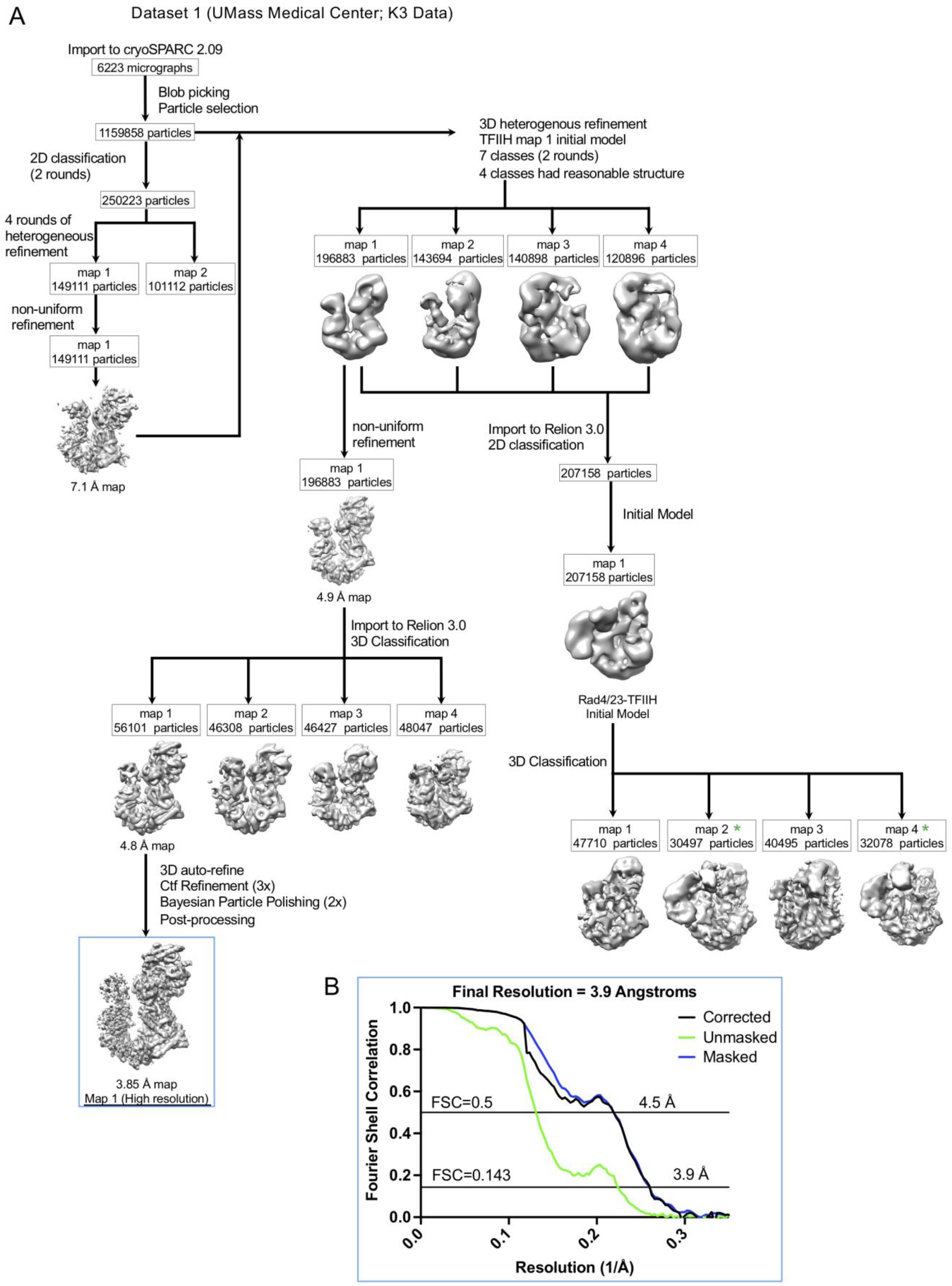
**(A)** Summary of data processing procedures and workflow of TFIIH/Rad4-23-33/DNA complex with ATP from K3 data collection. In short, data was motion corrected by MotionCorr2 and imported to cryoSPARC for CTF correction, particle picking and 2D classification. An initial reconstruction showed clear and novel TFIIH conformation at 4.9 angstroms. Import into Relion 3.0, followed by further 3D classification followed by CTF refinement, Bayesian polishing, and post processing yielded a reconstruction of 3.9 angstroms. **(B)** Fourier shell correlation (FSC) curves by postprocessing in high resolution TFIIH map (Map 1) in Relion 3.0 from K3 data. The resolution is estimated at FSC=0.143.

**Supplemental Figure 5.**
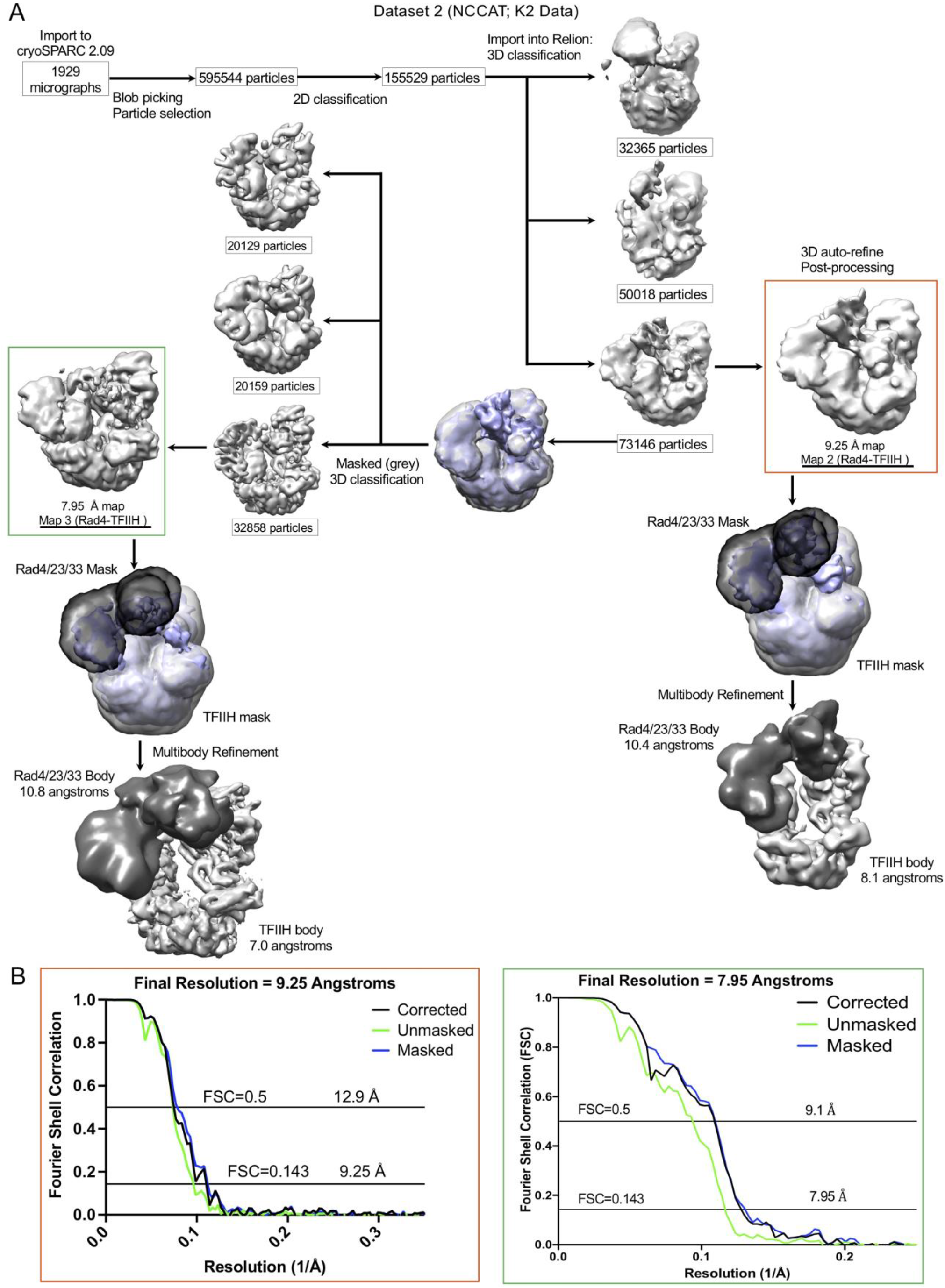
**(A)** Summary of data processing procedures and workflow of TFIIH/Rad4-23-33/DNA complex from K2 data collection. Similar to processing of K3 data (A) but focused on TFIIH/Rad4-23-33/DNA complex and conducted in Relion 3.1. **(B-C)** Fourier shell correlation (FSC) curves of Map 2 (B) and Map 3 (C).

**Supplemental Figure 6.**
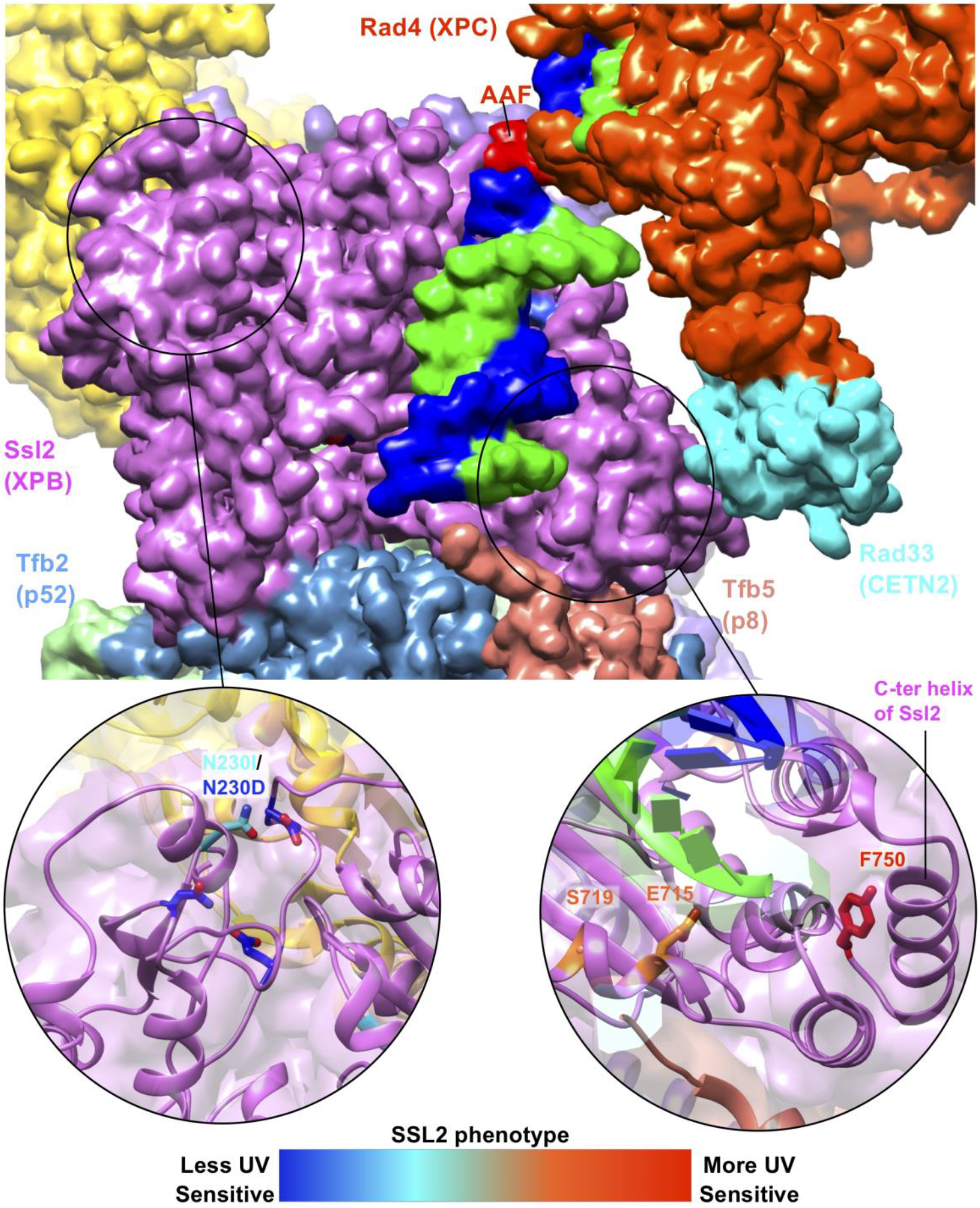
Mutations of Ssl2 at the interface with Rad33 and Tfb5 confer UV sensitivity. Mapping of mutations (red, severe; blue, mild) in Fig. 5D onto the structure of the TFIIH/Rad4-23-33/DNA complex.

**Supplemental Figure 7.**
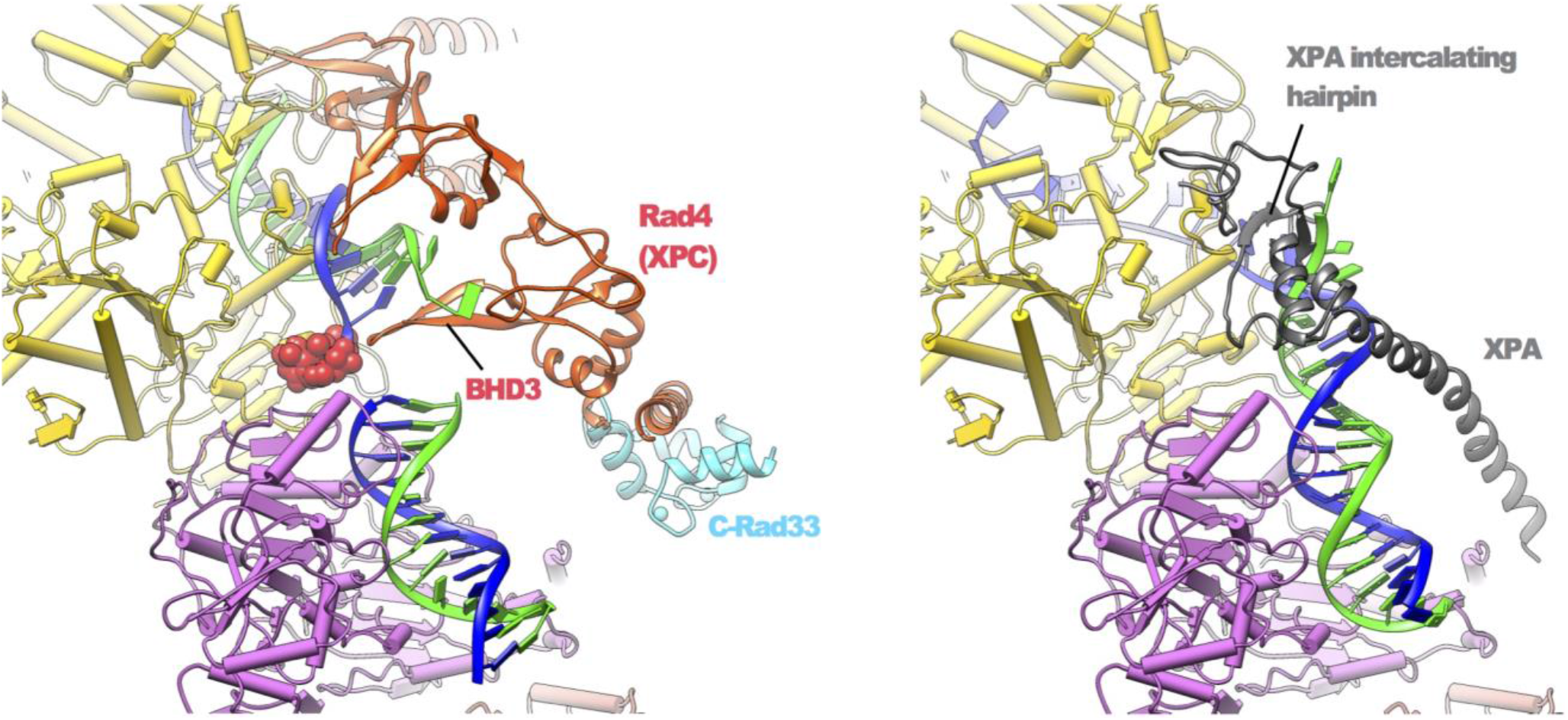
DNA binding sites of the yeast pre-unwound state (left) and previous structure of the human (artificially) unwound state (right, PDB:6RO4).

## Notes

### Competing Interest Statement

The authors have declared no competing interest.

